# A mutant fitness compendium in Bifidobacteria reveals molecular determinants of colonization and host-microbe interactions

**DOI:** 10.1101/2023.08.29.555234

**Authors:** Anthony L. Shiver, Jiawei Sun, Rebecca Culver, Arvie Violette, Charles Wynter, Marta Nieckarz, Samara Paula Mattiello, Prabhjot Kaur Sekhon, Lisa Friess, Hans K. Carlson, Daniel Wong, Steven Higginbottom, Meredith Weglarz, Weigao Wang, Benjamin D. Knapp, Emma Guiberson, Juan Sanchez, Po-Hsun Huang, Paulo A. Garcia, Cullen R. Buie, Benjamin Good, Brian DeFelice, Felipe Cava, Joy Scaria, Justin Sonnenburg, Douwe Van Sinderen, Adam M. Deutschbauer, Kerwyn Casey Huang

## Abstract

Bifidobacteria commonly represent a dominant constituent of human gut microbiomes during infancy, influencing nutrition, immune development, and resistance to infection. Despite interest as a probiotic therapy, predicting the nutritional requirements and health-promoting effects of Bifidobacteria is challenging due to major knowledge gaps. To overcome these deficiencies, we used large-scale genetics to create a compendium of mutant fitness in *Bifidobacterium breve* (*Bb*). We generated a high density, randomly barcoded transposon insertion pool in *Bb*, and used this pool to determine *Bb* fitness requirements during colonization of germ-free mice and chickens with multiple diets and in response to hundreds of *in vitro* perturbations. To enable mechanistic investigation, we constructed an ordered collection of insertion strains covering 1462 genes. We leveraged these tools to improve models of metabolic pathways, reveal unexpected host- and diet-specific requirements for colonization, and connect the production of immunomodulatory molecules to growth benefits. These resources will greatly reduce the barrier to future investigations of this important beneficial microbe.

## Introduction

*Bifidobacterium* species are early colonizers of the human gastrointestinal tract whose abundance has been associated with infant breastfeeding^1–4^. *Bifidobacterium* species are a focus of intense interest as probiotic interventions in early-life disorders associated with microbiome dysbiosis^5–7^. Specific associations between some *Bifidobacterium* species and health benefits include vitamin production^8^, immune training through production of small molecules such as indole-3-lactic acid^9–11^, stimulation of production of the anti-inflammatory signal IL-10^12^, and the production of other health-relevant small molecules such as conjugated linoleic acid^13^. However, many of these host-microbe interactions are species- or even strain-specific, making it imperative to understand their genetic and physiological basis before designing the next generation of *Bifidobacterium*-based interventions. Overcoming historical challenges in the genetic manipulation of Bifidobacteria^14^ is a necessary step toward that goal.

Systematic approaches connecting genotype to phenotype in bacteria have resulted in important insights into the physiology of many model organisms. An early effort mapped the phenotypic landscape of *Escherichia coli* by measuring colony size of >4,000 individual gene knockout strains in >100 unique growth conditions^15^. This study led to insight into multiple physiological aspects that extended even the vast knowledge base of *E. coli* molecular biology, and the chemical-genomic data set has subsequently served as a discovery platform for many laboratories using *E. coli* as a model system. Subsequent efforts to apply the same approach in other organisms have been successful in some ways but faced several challenges. In the limited number of bacteria for which targeted gene deletion is routine, the generation of genome-wide knockout libraries has proven expensive and time-consuming^16^. In the much larger number of bacteria for which gene knockout technology is non-existent or laborious, transposon insertion pools have enabled the interrogation of gene fitness across environments using insertion sequencing (INSeq) and variations like transposon sequencing (TnSeq)^17^. However, the laborious and costly nature of library preparation for INSeq/TnSeq have limited the scale of the resulting data sets.

Randomly barcoded transposons provide an opportunity to dramatically expand the scale of chemical-genomic screens by including a short molecular barcode near one inverted repeat of the transposon^18^. Once barcodes are associated with insertion sites via a single randomly barcoded transposon sequencing (RB-TnSeq) library, all subsequent assays of strain abundance can utilize the relatively simple barcode sequencing (BarSeq) library preparation. By reducing the time, cost, and effort of library preparation, randomly barcoded transposons enable large-scale chemical-genomic data sets to be generated for non-model organisms. This technology was deployed broadly in 32 bacteria and defined fitness effects of >11,000 genes with no previously known phenotypes^19^, but has yet to be applied broadly outside of the Pseudomonadota. Application of RB-TnSeq to *Bacteroides thetaiotaomicron* revealed the power of combining *in vitro* chemical-genomic data with fitness measurements during mouse colonization, demonstrating a *B. thetaiotaomicron* response to diet-dependent ammonium availability *in vivo*^20^ and providing a pathway toward dissecting the molecular mechanisms behind host colonization.

Incorporation of random barcodes into transposons has also benefited approaches for generating ordered mutant collections via isolation of single strains from random transposon pools by lowering the time, cost, effort, and complexity of identifying strains isolated in large progenitor collections. For *B. thetaiotaomicron*, we incorporated barcoding and flow cytometry into a protocol^21^ that enabled isolation of ∼2500 single insertion strains^22^. Such ordered mutant collections enable more detailed measurements of growth (lag phase, growth rate, cell death) and investigation of non-fitness related phenotypes such as enzyme activity in cell-free extracts or the production of metabolites^20^. Given that many of the health-promoting phenotypes of Bifidobacteria involve the production of small molecules, a comprehensive ordered library for the Bifidobacteria would be a powerful resource.

*Bifidobacterium breve* (*Bb*) is preferentially associated with infants in industrialized nations^2^. While *Bb* strains generally lack the capability to utilize the same breadth of human milk oligosaccharides (HMOs) as *Bifidobacterium infantis* strains, they can utilize some HMOs such as lacto-N-tetraose (LNT) and lacto-N-neotetraose (LNnT)^23^ and can cross-feed on monosaccharides released by other members of the gut microbiota^24,25^, including fucose. *Bb* strains produce indole-3-lactic acid^10^ and conjugated linoleic acid^13^, both of which have been associated with positive impacts on host health^26,27^. Importantly, previous work demonstrated that *Bb* strain UCC2003 could be transformed at sufficiently high efficiency to create a transposon-insertion library^28^. Thus, we focused on *Bb* UCC2003 for further study and development of genetic resources.

To create a compendium of mutant fitness in the Bifidobacteria, we generated a randomly barcoded transposon pool in *Bb* UCC2003. We used two complementary approaches to interrogate this transposon pool: growth *in vitro* in a chemical-genomic screen including 368 conditions, and colonization of two gnotobiotic hosts fed various diets. From this pool, we constructed an ordered single-transposon strain collection covering 1462 genes. We leveraged the compendium to curate more accurate and complete metabolic models, including pathways for the biosynthesis of amino acids, nucleotides, and vitamins and the breakdown of multiple carbohydrates. We used this information to identify a phenomenon of glucose toxicity that may shape the response of Bifidobacteria to changes in diet. We show that multiple biological processes contribute to cell shape regulation and that shape phenotypes are generally associated with fitness defects *in vitro* and *in vivo*, and we challenge the prevailing hypothesis that cell branching is a result of defects in cell wall synthesis. We demonstrate that Bifidobacteria have distinct stress responses from model organisms like *E. coli* when treated with genotoxic agents. Finally, we demonstrate that production of aromatic lactic acids in *Bb* contributes to growth by recycling redox cofactors NAD^+^ and NADPH.

### A compendium of mutant fitness in *Bifidobacterium breve* reveals genetic features of metabolism

We selected *Bifidobacterium breve* (*Bb*) UCC2003 as a model strain to expand genetic resources within the *Bifidobacterium* genus, due to its proven transformability^29,30^ and existing knowledge base. Building on optimized transformation protocols, we created a transposon-insertion pool consisting of >250,000 distinct mutants through the electroporation of *in vitro*–assembled transposomes (**Figure 1A, Figure S1, Table S1, S2**). Transposons in this library were barcoded to facilitate quantification of fitness using barcode sequencing (BarSeq)^31^.

**Figure 1:**
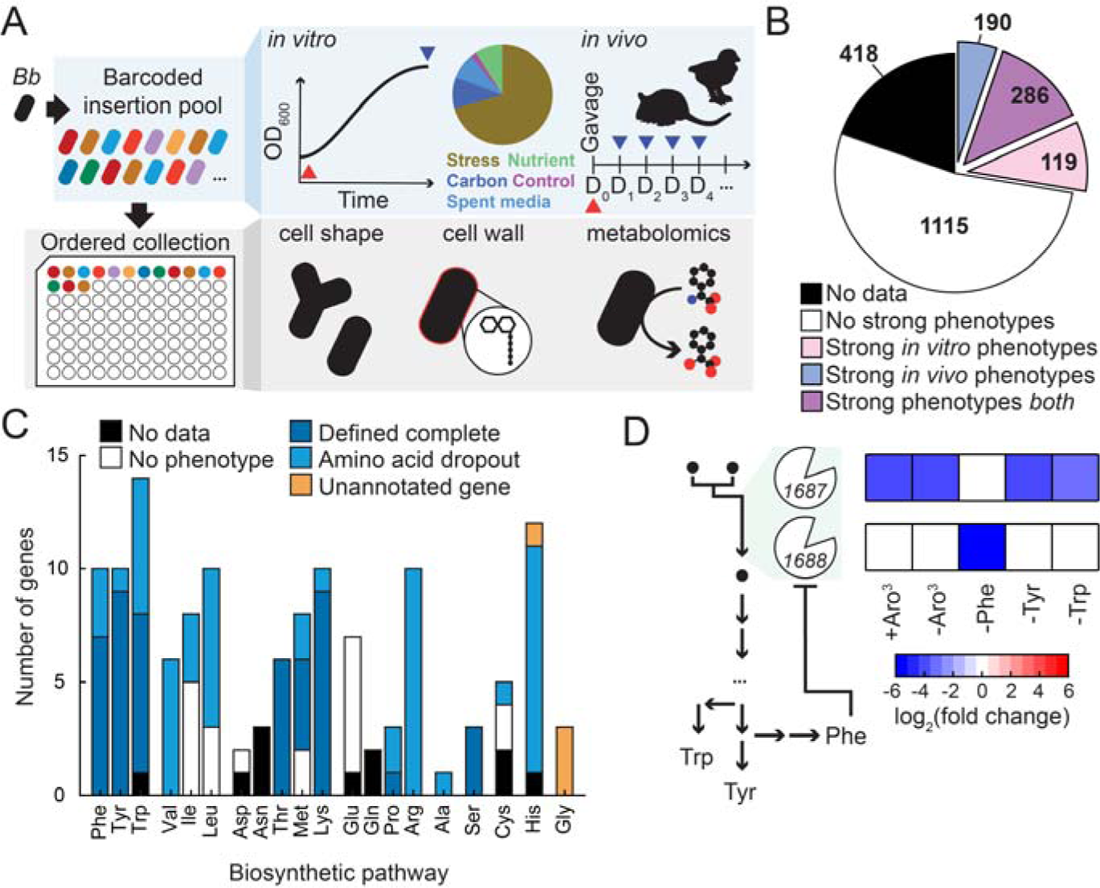
A mutant fitness compendium reveals genetic features of Bifidobacterial metabolism. A) Schematic of the workflow used to generate the fitness compendium. Top: randomly barcoded transposon insertion pools were screened with 368 perturbations *in vitro* and during colonization of germ-free hosts *in vivo*. Changes in relative abundance of strains between initial cultures (red triangle) and cultures post-perturbation (blue triangles) were estimated with barcode sequencing. A single time point after growth into stationary phase was sampled for each perturbation *in vitro*, while daily fecal samples (D_1_, D_2_, …) were collected *in vivo*. Bottom: a collection of 1462 insertion strains was isolated to enable interrogation of non-fitness phenotypes, including cell shape, cell wall architecture, and untargeted metabolomics. B) The fraction of the genome with at least one strong phenotype in the fitness compendium. Strong phenotypes were defined as having |fitness|>2 and |*t*|>5. 286/1710 genes had strong phenotypes both *in vitro* and *in vivo*, facilitating mechanistic investigation of colonization determinants. C) The fitness compendium informed models of amino acid biosynthetic pathways in *Bb*. Genes predicted to be involved in a pathway due to homology were grouped according to whether they had no data in the compendium (black), displayed no strong phenotypes (white), had growth defects in defined media with complete amino acids (dark blue), or had growth defects in defined media lacking the amino acid product of the pathway (light blue). Genes not predicted to participate in a pathway were assigned to it if the pathway had gaps and mutants had strong phenotypes in media missing the amino acid (orange). D) Details of biosynthetic regulation were revealed by the fitness compendium. Left: two genes, *Bbr_1687* and *Bbr_1688*, encode homologues of an enzyme predicted to catalyze the first step of the chorismate pathway: 3-deoxy-7-phosphoheptulonate synthase. Right: a heatmap of fitness phenotypes (log_2_–fold change in abundance) for both genes across multiple conditions. Mutants in *Bbr_1688* disrupt growth when phenylalanine is the only aromatic amino acid missing from the environment. Values are averages of at least two replicate experiments. Aro^3^: phenylalanine, tyrosine, tryptophan.

We used this transposon pool in three complementary investigations to generate a mutant fitness compendium in *Bb* (**Figure 1A**). First, we generated a chemical genomic dataset with fitness measurements in 368 conditions including antibiotics, nutrient dropouts, carbohydrate sources, and environmental variables relevant to host colonization such as temperature and osmolarity. Second, we colonized germ-free hosts (adult mice and chicks) on various diets to identify genes important for colonization. Third, we assembled an ordered collection of transposon insertion strains covering 1462 genes and measured individual growth curves and stationary-phase morphology of all strains in this collection across three growth conditions.

For sequencing-based measurements of mutant fitness, we relied on log_2_ fold-change in relative abundance (*fitness*) for effect size and a modified test statistic (*t*) for statistical significance. We defined cutoffs at |*fitness*|>0.5 and |*t*|>4 for significant phenotypes and |*fitness*|>2 and |*t*|>5 for strong phenotypes, as done previously^19^. The false discovery rate at both cutoffs was <5% (**Figure S2A**).

We verified the reliability of fitness measurements in the compendium in two ways. First, we inspected replicate measurements of mutant fitness, finding a high level of reproducibility both *in vitro* and *in vivo* (**Figure S2B-D**). Second, we evaluated the ability to identify known mutant fitness defects by examining growth on 7 carbohydrates that were the focus of previous studies^23,32–37^. We confirmed 10/12 known mutant phenotypes. Of the 2 genes without phenotypes, the first was *Bbr_0123* (*apuB*), which encodes an extracellular amylopullulanase that breaks starch down into shorter maltooligosaccharides for import^35^. The growth phenotypes of mutants in this gene may suppressed by cross-feeding in the transposon pool. The second was *Bbr_0010* (*lacZ2*), one of multiple β-galactosidases encoded in the *Bb* genome ^38^. The relative contribution of these redundant enzymes to total cellular β-galactosidase activity may depend on the exact growth conditions. In addition to recapitulating the majority of known mutant phenotypes, our compendium often expanded on prior findings, providing a more comprehensive picture of the uptake and breakdown of various carbohydrates (**Figure S2E**). For example, we identified a fitness phenotype for genes corresponding to 5/6 steps involved in the uptake and breakdown of fucose (**Figure S2E**). The lack of a growth phenotype for lactaldehyde reductase, the final step of the pathway, may reflect redundancy or the ability of *Bb* to excrete lactaldehyde as a waste product.

These positive outcomes motivated a broader survey of mutant fitness effects in the data set. Of the 1710 genes included in the fitness compendium, 595 displayed at least one strong fitness phenotype in at least one experiment, including measurements during host colonization and growth *in vitro* (**Figure 1B**). Of note, 48% of responsive genes (286/595) had phenotypes during both colonization and growth *in vitro*, an overlap which has been helpful previously for deriving mechanistic insights into host colonization^20^ and which we explore further to provide a global view of gene function in *Bb*.

We first focused on amino acid biosynthesis because the pathways have not been extensively studied in Bifidobacteria. Using data from *in vitro* experiments in which we left individual amino acids out of a chemically defined medium (**Table S3**), we curated models of biosynthetic pathways for all 20 amino acids. As part of the curation process, the compendium provided experimental evidence for the assignment of genes to 72 out of 89 reactions of amino acid biosynthesis (**Figure 1C**). Of the 17 steps without experimental evidence, 6 are essential and the remaining 11 may have redundant enzymes or pathways. Genes in the biosynthetic pathways for serine, threonine, lysine, and tyrosine displayed strong fitness defects in chemically defined media even with their respective amino acids present, suggesting that uptake of these amino acids is limited under the conditions tested (**Figure S2**). In some cases, the dataset provided evidence for enzyme regulation. *Bb* encodes two isozymes of the enzyme 3-deoxy-7-phosphoheptulonate synthase, *Bbr_1687* and *Bbr_1688*, which catalyze the first committed step of aromatic amino acid biosynthesis. Distinct phenotypes between these genes revealed that one of the two isozymes (*Bbr_1688*) is likely responsive to phenylalanine levels, potentially through feedback regulation^39^ (**Figure 1D**). The data also connected novel genes to gaps in pathways for which there were no homology-based predictions (**Figure 1C**).

### Discovery of Bifidobacterial enzymes involved in histidine biosynthesis

Although candidate genes responsible for many steps of amino acid biosynthesis in *Bb* could be identified based on conservation, we were unable to computationally annotate the full pathways for histidine and glycine^40^. However, *Bb* was able to grow in media lacking either amino acid and chemical sensitivities from the fitness compendium revealed enzymes that could fill the apparent gaps in these pathways. For histidine biosynthesis, the penultimate enzyme, histidinol phosphate phosphatase, was missing from an otherwise complete pathway (**Figure 2A**). We focused on histidine biosynthesis and characterized the putative histidinol phosphate phosphatase further.

**Figure 2:**
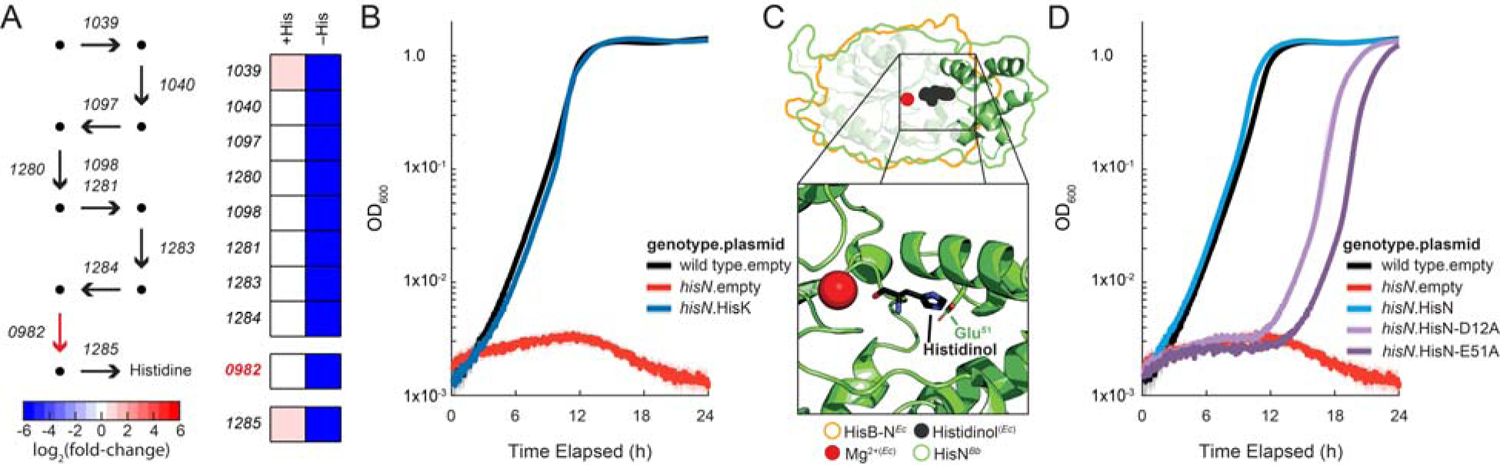
Identification of a Bifidobacterial histidinol phosphate phosphatase. A) Left: a sequence homology–based model of histidine biosynthesis identified candidate enzymes with strong phenotypes in defined media lacking histidine for every step of the pathway except histidinol phosphate phosphatase (red arrow). Right: an enzyme in the HAD family of phosphatases, *Bbr_0982*, was investigated as a candidate enzyme because of its shared phenotypes with the rest of the pathway. Points in the pathway represent pathway intermediates, and arrows represent enzymatic steps. The fitness phenotypes (log_2_–fold changes in abundance) are shown as a heatmap. Values in the heatmap are averages of at least two replicate experiments. B) Heterologous complementation confirmed that HisN (encoded by *Bbr_0982*) is a histidinol phosphate phosphatase. The OD_600_ (optical density at λ=600 nm) is plotted on a logarithmic scale. The mean of replicate growth curves (*n*=5) is shown as a solid line while the transparent region represents the 95% confidence interval (1.96×standard error). An insertion strain in *Bbr_0982* (*hisN*) was unable to grow in chemically defined media without histidine. Heterologous expression of histidinol phosphate phosphatase from *Bacillus subtilis* (HisK) rescued growth in a mutant background. C) Alignment of an AlphaFold structural prediction for HisN*^Bb^* (green outline, green cartoon structure) with HisB-N*^Ec^* (orange outline), the histidinol phosphate phosphatase subunit from *E. coli* (PDB ID: 2FPU), revealed a conserved domain in HisN*^Bb^* with a similar structure (transparent cartoon) to HisB-N and a domain unique to Bifidobacteria (opaque cartoon). The non-conserved domain of HisN*^Bb^* included residue Glu-51, which the alignment positioned to form an electrostatic interaction with the imidazole nitrogen of histidinol (inset, bottom). Mutations disrupt the function of HisN. The OD_600_ (optical density at λ=600 nm) is plotted on a logarithmic scale. The mean of replicate growth curves (*n*=5) is shown as a solid line and the transparent region represents the 95% confidence interval (1.96×standard error). Ectopic expression of *Bb* HisN rescued growth in chemically defined media lacking histidine. However, mutations in the conserved active site (HisN-D12A) and non-conserved domain (HisN-E51A) were impaired in their ability to complement growth.

Insertions in a predicted haloacid dehalogenase (HAD)–family phosphatase, *Bbr_0982* (*hisN*), had strong and specific fitness defects in media without histidine, suggesting that the gene encodes the missing histidinol phosphate phosphatase (**Figure 2A**). Supporting this functional assignment, the growth phenotype of *hisN* was complemented by expression of the native gene (*hisN^Bb^*) or heterologous expression of histidinol phosphate phosphatase from *B. subtilis* (*hisK^Bs^*) (**Figure 2B**).

As a member of the HAD family of phosphatases, HisN*^Bb^*is a distant homologue of the histidinol phosphate phosphatase domain in the bifunctional enzyme HisB from *E. coli* (HisB-N*^Ec^*). Members of this family share a highly conserved N-terminus, which encodes the active site residues that chelate the catalytic magnesium cation. However, sub-families differ at a variable insertion domain that confers substrate specificity to the enzyme^41^. The variable domain of HisN*^Bb^* is not conserved outside the Bifidobacteria, explaining the inability of a sequence-based algorithm to confidently predict its specific function. However, a structural prediction of HisN*^Bb^*using Alphafold^42^ aligned closely to a crystal structure of HisB-N*^Ec^*at the conserved domain^43^, and placed the carboxyl group of glutamate at residue 51 of HisN*^Bb^* in position to interact with the nitrogen of the imidazole ring of histidinol phosphate (**Figure 2C**). To test this prediction, we measured the ability of HisN-D12A (conserved domain, catalytic mutant) and HisN-E51A (variable domain, substrate binding mutant) to complement the *hisN* mutant background. Both mutants exhibited slow growth in histidine-free media (**Figure 2D**), supporting the structural prediction of the HisN^Bb^ non-conserved domain. These results demonstrate the power of a fitness compendium to associate novel genes with biological processes, independent of sequence homology.

### A high-glucose diet is toxic to *Bb* during mono-colonization

We next sought to determine if our dataset could reveal metabolic requirements in more complex conditions like host colonization. We gavaged the *Bb* transposon pool into adult mice fed one of three diets: a standard diet (SD), a high-sucrose and high– saturated fatty acid dietary mix referred to as a Western diet (WD), and a diet deficient in microbially accessible carbohydrates (MD) in which the sole carbohydrate supplied was glucose. We also fed the *Bb* transposon pool to newly hatched germ-free chicks fed a standard chick diet (CD). The relative abundance of strains was monitored by sampling feces over the course of a month.

Strikingly, mutants in genes associated with sugar uptake expanded to dominate all four gut environments across both hosts within the first week of colonization (**Figure 3A**). This expansion included a multi-gene phosphotransferase system (PTS) that dominated SD-fed mice (*Bbr_1892–1894*) and genes encoding an ABC transporter (*Bbr_1844*, *Bbr_1845*, and *Bbr_1847*) that took over the pool in WD-fed mice and CD-fed chicks. In MD-fed mice, sugar uptake mutants expanded to represent the majority of the transposon pool within a single day after gavage. These mutants included a single-gene PTS (*Bbr_1594*) as well as the common phospho-relay genes (*Bbr_0183* and *Bbr_0182*) and a predicted transcriptional regulator of the PTS (*Bbr_1593*).

**Figure 3:**
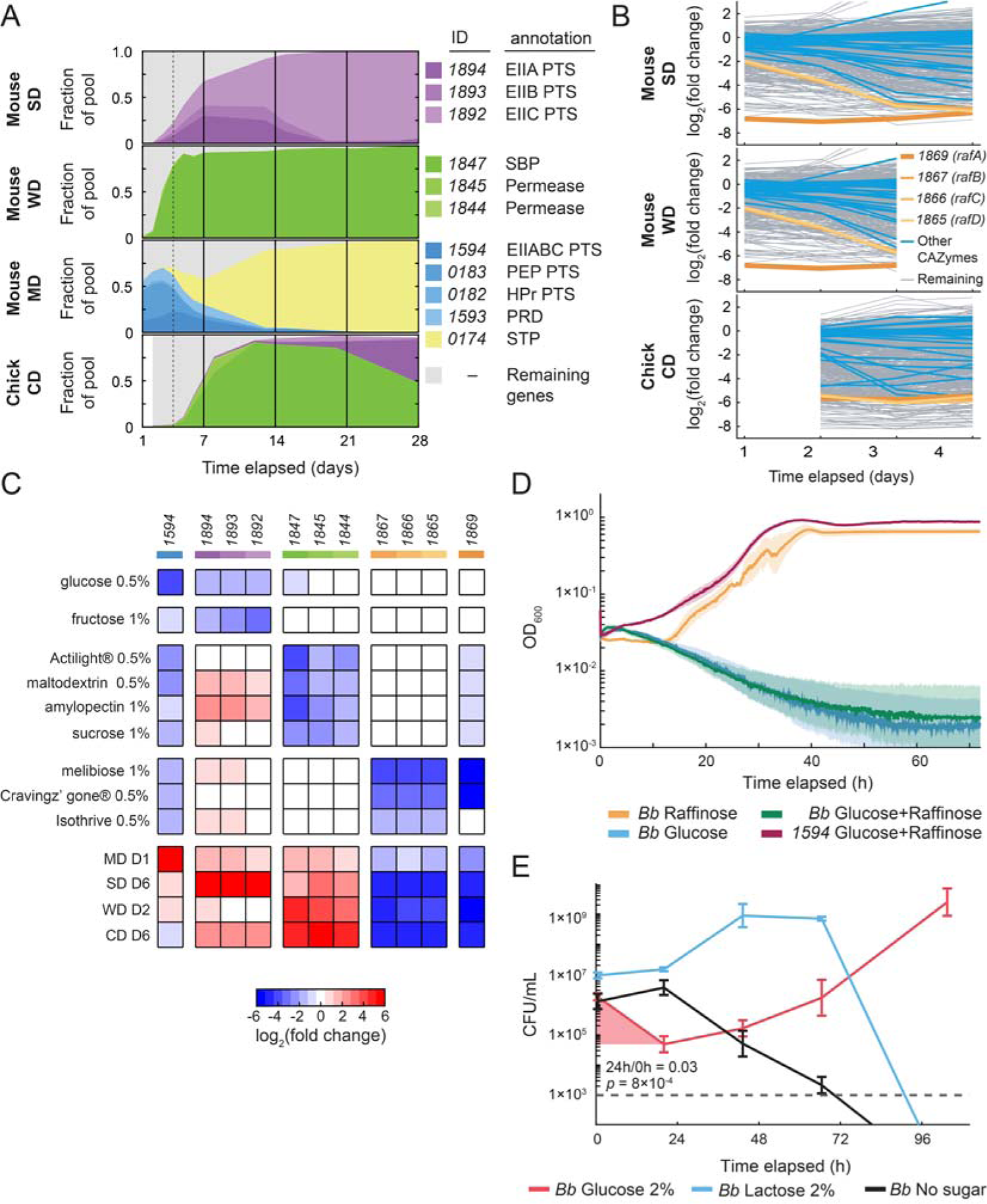
Selective pressures during colonization reveal a glucose toxicity phenomenon in *Bb*. A) Across every combination of host and diet, the transposon pool was dominated by a small subset of mutants in sugar transporters within one week. Reads from all insertions that map within a given gene are plotted as a fraction of the total pool and as a function of time elapsed post-gavage. Data from cage mates were averaged at each time point. Plots from four combinations of host and diet are shown: female adult mice on a standard mouse diet (SD, top), female adult mice on a high-sucrose/high-polyunsaturated fat “Western” diet (WD, second from top), female adult mice on a microbially accessible carbohydrate–deficient diet (MD, second from bottom), and newborn chicks on a standard chicken diet (CD, bottom). The transposon pool in the SD-fed mice was dominated by mutants in a multi-gene EII complex of a phosphotransferase system (PTS, purple, *Bbr_1894*–*1892*). In the WD-fed mice, mutants in an ABC transporter dominated (green, *Bbr_1844*,*1845*,*1847*). The same mutants dominated the CD-fed chicks (bottom). The transposon pool in the MD-fed mice was dominated within one day by mutants in a single-gene EII complex of a PTS (*Bbr_1594*), as well as the universal upstream genes of the PTS pathway (blue, *Bbr_0182,0183*) and a predicted regulator (*Bbr_1593*). Early time points up to day 4 (dashed vertical line) were investigated for fitness determinants; after this time, biases in the pool made fitness phenotype calculations less reliable. B) One of the most important carbohydrate-active enzymes (CAZymes) for colonization, regardless of host or diet, was *Bbr_1869* (*rafA*). The fitness values (log_2_–fold change in abundance) of genes are plotted for the first four days of the SD-, WD- and CD-fed colonization experiments. The fitness defect of *rafA* (dark orange) was the strongest among defect among all CAZymes (blue). In mice, *rafA* colonization defects were the strongest among all genes in the pool (grey). Mutations in the transporter associated with *rafA* (*Bbr_1867–1865* (*rafBCD*), shades of orange) were also deleterious for colonization in all hosts and diets. C) Phenotypes in the mutant fitness compendium connect transporters to their substrates. A heatmap of fitness values is shown for genes encoding the three transporters that take over during colonization (blue, purple, and green) and the transporter associated with *rafA* (orange) that is important for colonization in all hosts and diets. D) Glucose prevents growth in media with low amino acid content. Growth curves are plotted for wild-type cultures with 1.5% raffinose (orange), 0.15% glucose (blue), and 1.5% raffinose+0.15% glucose (green) and for *Bbr_1594* with 1.5% raffinose+0.15% glucose (burgundy). The growth medium is CDM2 lacking all amino acids except cysteine and phenylalanine. The mean of individual measurements (*n*=3) is plotted as a solid line and the 95% CI (1.96×SE) is plotted as a patch. Growth of wild type, but not the transport mutant (*Bbr_1594*), is inhibited by glucose, even as a supplement to other carbohydrates. 95% CI: 95% confidence interval, SE: standard error. E) Glucose kills *Bb* in amino acid–poor environments. Colony forming units (CFU) of wild type were enumerated after transfer to CDM2 lacking all amino acids except cysteine and phenylalanine. The carbon source was 2% lactose (blue), 2% glucose (red), or no carbohydrate (black). Error bars represent the 95% CI (1.96×SE, *n*=4). The fraction of viable cell counts of *Bb* dropped to 0.03 (95% CI: 0.01–0.06, *p*=8×10^-4^) in the first 24 h after transfer to media with glucose, showing that glucose uptake is toxic under these conditions. The dotted grey line marks the lower limit of detection in this assay.

Despite the strong fitness advantage of these transporter mutants, we were able to quantify the abundance of other barcodes in the first days of colonization, providing an early window into the fitness determinants of colonization in *Bb*. Examining the carbohydrate active enzymes (CAZymes), we found that mutations in *Bbr_1869* (*rafA*) exhibited some of the strongest fitness defects across host and diet (**Figure 3B**). These defects were especially strong in mice, for which *rafA* was the only CAZyme with strong fitness effects on the first day after gavage. The enzyme RafA has been experimentally verified as an α-galactosidase important for growth on raffinose-family oligosaccharides (RFOs) like raffinose, melibiose, and stachyose^33^. Mutations in the ABC transporter associated with *rafA*, *Bbr_1865–1867* (*rafB-D*), also displayed strong colonization defects across host and diet (**Figure 3B**). The RFOs raffinose and stachyose are detected in cecal and fecal contents of germ-free mice^44^, consistent with RFOs being an important carbon source for *Bb* during colonization.

Having identified sugar transporters with positive and negative fitness effects during colonization, we turned to our mutant fitness compendium and ordered collection to identify potential substrates of these transporters. We demonstrated glucose as the major substrate for *Bbr_1594*, fructose as the substrate for *Bbr_1892–1894*, and multiple substrates, including several prebiotics, for the two ABC transporters (**Figure 3C, Figure S3A**). The substrates identified for the two PTS transporters were somewhat unexpected, given that *Bbr_1594* had previously been proposed to be a fructose transporter based on its phenotype during heterologous expression in *E. coli*^45^. However, both PTSs had significant growth defects on both glucose and fructose, and mutations in the respective PTS genes did not completely eliminate growth on their expected substrates (**Figure S3A**), suggesting cross-specificity.

Identifying carbohydrate substrates of the sugar transporters revealed an unexpected and counter-intuitive connection to their positive fitness effects during colonization. In diets supplemented with purified sugars (glucose in MD and sucrose/starch in WD), a selective pressure in the *Bb* transposon pool apparently existed to eliminate transport of the added sugars (**Figure 3C**). To explore this phenomenon further, we sought to recreate the selective pressure for transporter loss *in vitro*. We first focused on co-utilization of raffinose with model substrates of the transporters that swept the transposon pool *in vivo*. We found that addition of glucose had progressively negative impacts on growth as the amino acid content of the growth medium was lowered (**Figure S3B**). In defined media containing only L-cysteine and L-phenylalanine, even a relatively small amount of glucose (0.15% w/v) was sufficient to halt the growth of *Bb* (**Figure 3D**). Mutation of the glucose transporter rescued growth under these conditions (**Figure 3D**), recapitulating the selective pressure originally detected in mice fed the MD.

The negative effect of glucose was not specific to co-utilization with raffinose, as the *Bbr_1594* mutant rescued growth even when glucose was the only carbohydrate added (**Figure S3C**). This finding led us to hypothesize that glucose was toxic to *Bb* in an amino acid–poor environment. To test this hypothesis, we enumerated colony forming units (CFUs) of *Bb* in amino acid–poor defined media supplemented with glucose, lactose, or no carbohydrate. As predicted, glucose was bactericidal for *Bb* under these conditions, eliminating >95% of the initial CFUs within 24 h and requiring 76 h for recovery to initial levels (**Figure 3E**).

While the environment that *Bb* faces during colonization is unlikely to exactly match the *in vitro* growth conditions that enhance glucose toxicity, we hypothesize that *Bb* reaches an analogous physiological state during colonization that is sensitized to glucose and generates the strong selective pressures inferred for mice fed an MD. Given the unique toxicity of glucose *in vitro*, and the different time scales of selection *in vivo* across diets, we expect that the selective pressure acting against the other transporters is distinct from glucose in the MD.

### A Bifidobacterial protein contributes to survival of off-target effects of genotoxic stress

We were struck the similarity of early fitness requirements for colonization of adult mice and chicks on their respective standard diets (*r*=0.68, *p*=3⨉10^-232^, 95% CI=0.65–0.70, day 2 of colonization) (**Figure 4A**) despite differences in host organism, diet, and colonization method (**Methods**). However, notable differences existed between the two experiments. Colonization of chicks exposed more than double the number of genes with strong fitness defects than adult mice; 186 and 89, respectively (**Figure 4A**). We found an enrichment in chicks for genes in the biosynthesis of lysine and threonine (*NES=*-1.46 *fdr=*0.22 and *NES*=-1.39 *fdr*=0.18, respectively) (**Figure 4A**). Surprisingly, we also found an enrichment of genes involved in homologous recombination repair (HRR) (NES=-1.34 fdr=0.31, AdnAB pathway) (**Figure 4A**). An uncharacterized gene, *Bbr_0059* (*scp3*), was also specifically required during chick colonization (**Figure 4A**). We found that *scp3* was sensitized to multiple genotoxic agents *in vitro*, causing it to group with genes associated with HRR and nucleotide excision repair (NER) in hierarchical clustering of the mutant fitness compendium (**Figure 4B**). Given its importance for survival of genotoxic agents and its specific connection to chick colonization, we sought to explore the biological function of Scp3 further.

**Figure 4:**
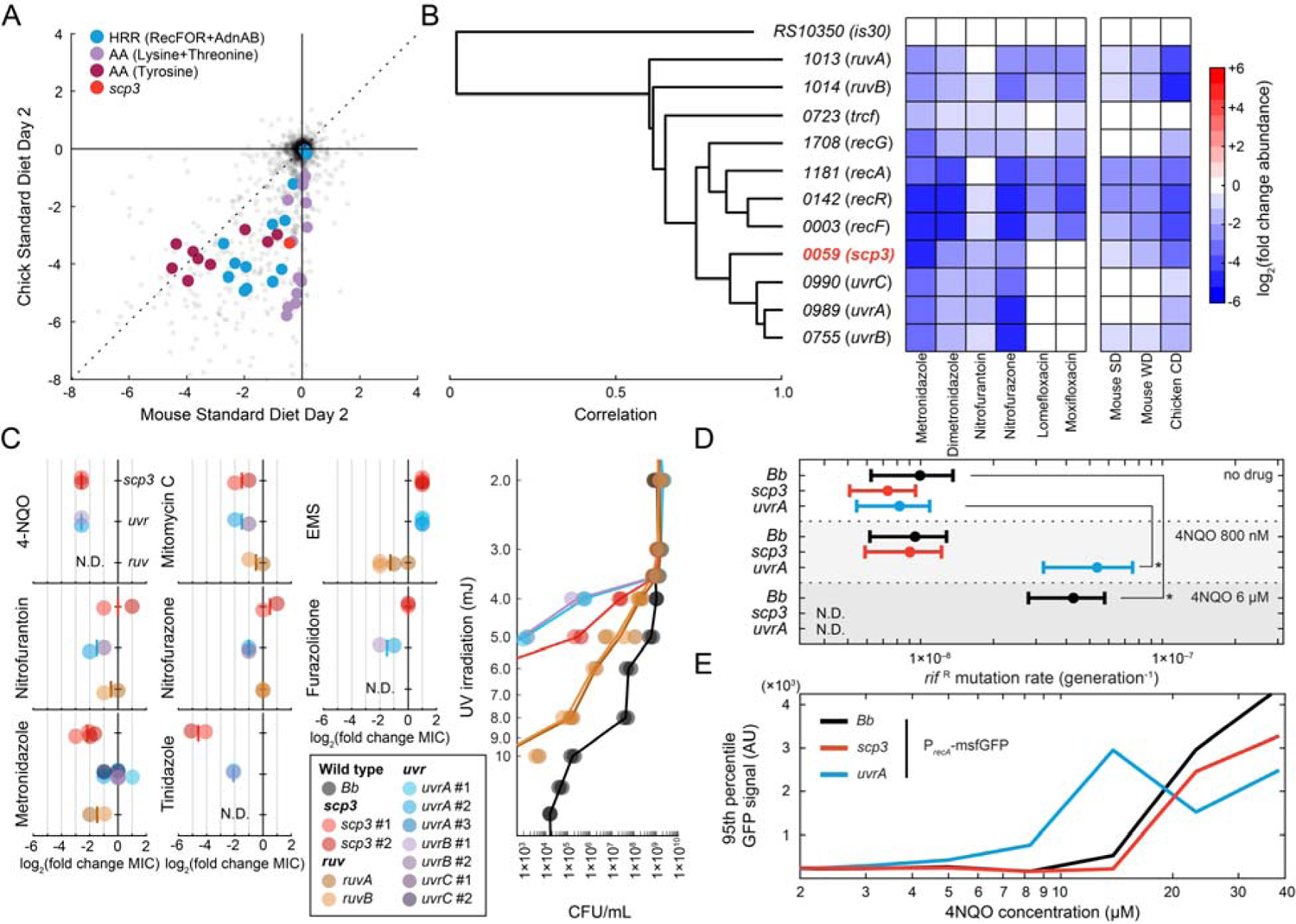
A Bifidobacterial protein contributes to survival of off-target effects of genotoxic stress. A) Some fitness requirements differed between colonization of mice and chicks, each fed their standard diets. Tyrosine biosynthesis was important for colonization in both conditions, while threonine and lysine biosynthesis were primarily important for colonization of chicks. Genes predicted to contribute to homologous recombinational repair (HRR) were more important for colonization of chicks than mice on a standard diet. A gene of unknown function, *Bbr_0059* (*scp3*), was also important for colonization of chicks but not mice. B) Hierarchical clustering of the mutant fitness compendium grouped *scp3* with genes responsible for nucleotide excision repair (NER, *uvrABC*) and homologous recombinational repair (HRR, *recAFRG*). Like mutants in the NER pathway, *scp3* was not sensitive to the fluoroquinolone antibiotic family. C) *scp3* mutants exhibited overlapping but distinct sensitivities to mutants of NER and HRR. The change in minimum inhibitory concentration (MIC) for all genotoxic agents except ultraviolet radiation (UV) was quantified for *scp3*, *uvrAB* (NER), and *ruvAB* (HRR) using a liquid broth dilution approach. MIC changes are plotted as log_2_–fold change in MIC compared to a wild-type control. Top row: *scp3* and *uvr* have similar sensitivities to antibiotics traditionally associated with NER (4-nitroquinoline 1-oxide (4-NQO), mitomycin C) and do not contribute to resistance to the alkylating agent ethyl methylsulfonate (EMS). Middle row: *uvr* mutants are more sensitized than *scp3* to antibiotics in the nitrofuran family (nitrofurantoin, nitrofurazone, and furazolidone). Bottom row: *scp3* is more sensitized than *uvr* to antibiotics in the 5-nitroimidazole family (metronidazole and tinidazole). Far right: sensitivity to UV irradiation was quantified using spot dilutions on an MRS agar surface. CFU/mL are plotted as a function of UV irradiation (mJ). Both *scp3* and *uvr* mutants are sensitized to UV. D) 4NQO-induced mutagenesis rate was unaffected in an *scp3* background. A fluctuation test was performed to estimate 4NQO-induced mutagenesis rates in wild type, *scp3*, and *uvrA*. The rifampicin resistance mutagenesis rate is plotted for these strains at 0, 800 nM, and 6 µM 4NQO. Error bars represent the 95% CI. *scp3* and *uvrA* lowered the MIC of 4NQO to a similar degree (C), but only the *uvrA* mutation exhibited an increase in mutagenesis rate at sub-MIC concentrations. Data were not collected for the mutant strains at the highest concentration of 4NQO (N.D.) because the concentration was lethal for these strains. *: *p*<0.05, 95% CI: 95% confidence interval. E) Induction of the SOS response was unaffected in an *scp3* background. The 5’ UTR of *recA*, including the promoter and SOS box, was fused to *msfGFP* on a plasmid to create an SOS reporter. Plotted is the 95^th^ percentile of single-cell GFP signal (median value, background subtracted) in wild type, *uvrA*, and *scp3* as a function of 4NQO concentration. The concentration of 4NQO necessary to induce the SOS response was lower in the *uvrA* background, but not *scp3*.

While co-clustering of *scp3* mutants with DNA damage repair pathways suggested similar roles in the cell, many genotoxic agents have cellular targets other than nucleic acids^46–56^. Therefore, as a first step toward identifying a molecular function of Scp3, we tested the hypothesis that it has a direct role in DNA damage repair. We determined changes in minimum inhibitory concentration (MIC) for an expanded set of genotoxic agents, testing for differences between *scp3* and genes in NER (*uvrA*/*uvrB*) and HRR (*ruvA*/*ruvB*). Mutants in all pathways were sensitized to compounds classically associated with NER, including mitomycin C, 4-nitroquilone-1-oxide (4NQO), and UV exposure (**Figure 4C**). However, *scp3* and *uvrA*/*uvrB* could be distinguished by their relative sensitivities to the 5-nitroimidazole and nitrofuran families, respectively (**Figure 4C**). Mutations in *ruvA*/*ruvB* were unique in their sensitivity to the alkylating agent ethyl methanesulfonate (**Figure 4C**).

The observation that mutants in the HRR and NER pathways have susceptibilities not shared by *scp3* indicated that Scp3 does not play an essential role in either repair pathway but did not exclude a distinct role of Scp3 in DNA repair. To test this possibility, we tested for a genetic interaction between *scp3* and two known DNA repair pathways, translesion synthesis and SOS, using 4NQO as a representative stressor. We quantified induced mutagenesis rates in wild type, *scp3*, and *uvrA* by enumerating rifampicin-resistant mutant colonies after exposure to 4NQO. 4NQO concentrations that were inhibitory to growth for *scp3* and *uvrA*, but sub-inhibitory for wild type, elevated the mutagenesis rate of wild-type cultures (**Figure 4D**). At a lower concentration of 4NQO that was sub-inhibitory for *scp3* and *uvrA* mutants, only the *uvrA* mutant had a significantly elevated mutagenesis rate relative to wild type (**Figure 4D**). Neither *scp3* nor *uvrA* altered the mutagenesis rate in the absence of 4NQO (**Figure 4D**). These results indicate that loss of function of *uvrA*, but not *scp3*, causes the accumulation of genetic lesions that require mutagenic DNA polymerases at lower concentrations of 4NQO.

We next used a fluorescent reporter to measure activation of the SOS response in response to 4NQO. We fused the 5’ UTR of *recA* to msfGFP on a plasmid and used it to transform wild type, *scp3*, and *uvrA*. In a wild-type background, fluorescence increased dramatically in a subset of cells at 4NQO concentrations near the MIC (**Figure 4E**), indicating that the construct was functional as an SOS reporter. In a *uvrA* background, the SOS response was induced at a lower concentration of 4NQO compared to wild type, consistent with the lower MIC for 4NQO in the NER^-^ background (**Figure 4E**). In contrast to *uvrA*, the SOS response in *scp3* background tracked with that of wild-type cells (**Figure 4E**). These results indicate that loss of *uvrA* function, but not *scp3*, leads to double-stranded DNA breaks at a lower concentration of 4NQO.

The *scp3* gene is as important for *Bb* survival of genotoxic agents as genes in traditional DNA repair pathways, but multiple lines of evidence suggest that its molecular function is independent of DNA repair. Distinct antibiotic sensitivities demonstrate that Scp3 is not an essential component of either NER or HRR in *Bb* (**Figure 4B, C**). Loss of *scp3* function in the face of genotoxic stress neither increases mutagenic DNA repair (**Figure 4D**) nor induces the SOS-response (**Figure 4E**). Many genotoxic stressors like 4NQO are thought to have DNA as their primary cellular target^57–59^, in part because of the magnitude of the sensitization in mutants of DNA repair pathways^57,59^. While other cellular damage is known to occur, it remains relatively unexplored as a mechanism of killing by antibiotics. Our results suggest that Scp3 is a stress response protein that counteracts this damage. Further work to determine the molecular function of Scp3 may reveal not only an unappreciated mode of antibiotic action in the gut microbiome but also the unique colonization requirements of chicks.

### Bifid cell shape is associated with poor growth and host colonization

Bifidobacteria were first named for their propensity to form Y-shaped (bifid) cells when cultured from the stool of newborn babies^60^. Bifid cell shapes are often only present at low frequencies when Bifidobacteria are visualized directly from stool, but can become prevalent after subculturing, especially in nutrient-poor media^61–63^. Despite the prominence of bifid cell shapes in our conception of the Bifidobacteria, the physiological role and mechanistic basis of branching remain controversial. A long-standing model ^64^ proposes that branching occurs due to defects in cell wall synthesis. This model is supported by the observation that a strain of *Bifidobacterium bifidum* whose growth is dependent on the cell wall precursor N-acetyl-glucosamine formed bifid and bulged cell shapes when N-acetyl-glucosamine was omitted from the growth medium^65^. However, while starvation for cell wall precursors may be sufficient to promote bifid cell shape, whether it is necessary remains an open question. Recognizing the potential of our ordered collection for exploring the genetic determinants of Bifidobacterial cell shape, we set out to characterize abnormal shapes within the library and connect bifid cell shape to growth and changes in cell wall architecture.

We first characterized the chemical structure of the peptidoglycan cell wall and shape phenotypes of wild-type cells as a reference for the genome-scale ordered collection. After developing a UPLC-MS protocol for quantifying the chemical composition of *Bb* peptidoglycan (**Methods**), we detected the expected pentapeptide of *Bb*^66,67^ along with various crosslinked species (**Figure 5A**, **Table S4**). Furthermore, we determined that two prominent features of the *Bb* stem peptide, amidation of D-glutamate and addition of the glycine crossbridge, occur after the formation of the lipid II precursor (**Figure S5A**). We next characterized the propensity of *Bb* to form bifid and other non-rod-like shapes. Wild-type cells were largely rod shaped in nutritionally complete rich medium (MRS), and mutants in the transposon insertion pool were largely rod shaped in the colon of chicks a week after colonization (**Figure 5B**). However, bulged and branched shapes of wild-type cells were more frequent in a sub-optimal rich medium (MRS*) and a defined medium lacking most amino acids (CDM2-minimal AA) (**Figure 5B**). These data confirmed that *Bb* can serve as a representative model system for Bifidobacterial cell shape and its connection to cell wall architecture.

**Figure 5:**
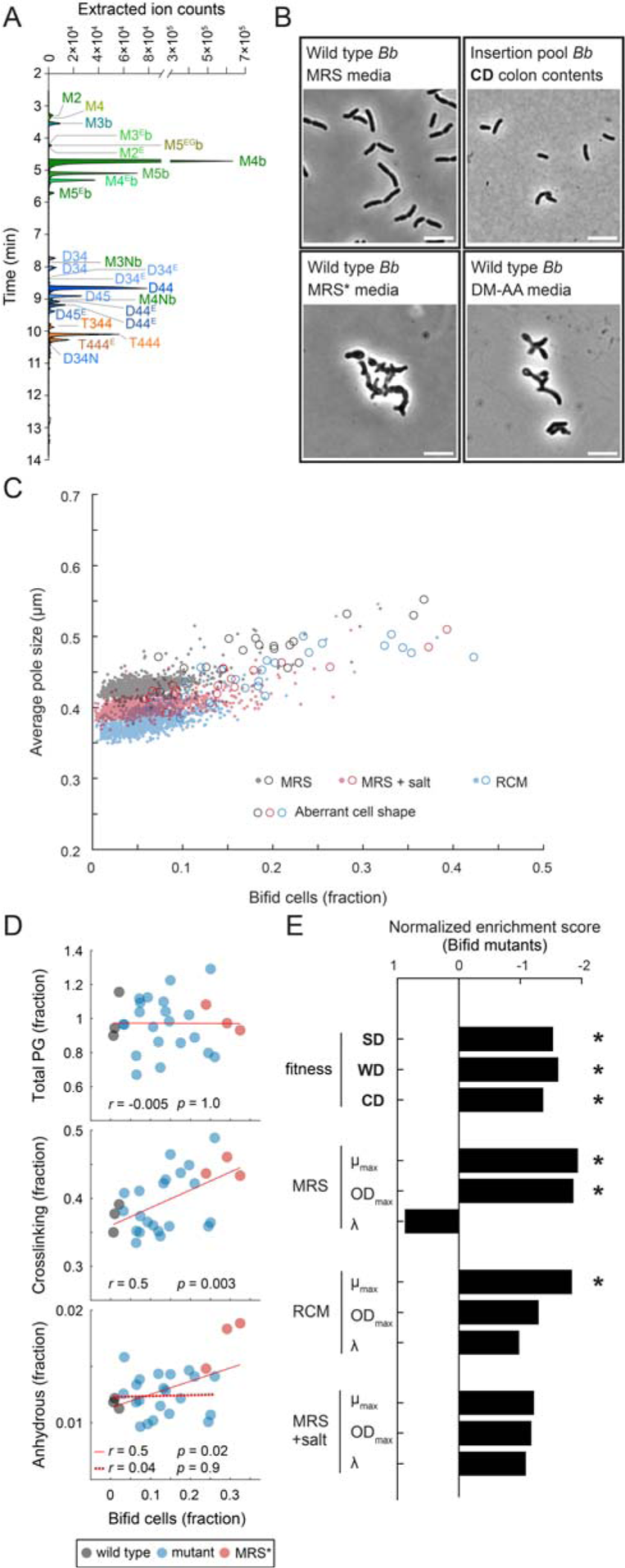
Genetic determinants of bifid cell shape reveal pervasive fitness defects. A) Extracted ion profile of *Bb* muropeptides. The extracted ion count of muropeptides is plotted as a function of UPLC retention time. Peaks identified by MS are labeled on the plot. Monomers (M, green), dimers (D, blue), and trimers (T, orange) are color coded. The exact chemical structure of detected muropeptides is recorded in **Table S4**. B) Bifid cell shape is influenced by nutritional state in *Bb*. *Bb* adopts a normal rod-like shape in the rich medium MRS (top left) and during host colonization (top right). Aberrant shapes such as bulges and branches were more abundant in nutritionally deficient media like MRS without meat extract (MRS*, bottom left) and a chemically defined medium lacking all amino acids except cysteine (bottom right). C) A screen for cell shape defects identified multiple genes that influence *Bb* morphology. Shown are quantification of two cell shape parameters, bifid cell abundance and pole width (Methods), for stationary-phase cells in three media: MRS, MRS with 300 mM NaCl, and RCM. A general trend connects pole width to extent of branching across the population. Outliers in cell shape that were confirmed by visual inspection to have any aberrant shapes (**Table S5**), even beyond pole size and branching, are shown as open circles. D) Bifid cell abundance was not correlated with defects in cell wall assembly. Bifid cell abundance in stationary phase is plotted against total peptidoglycan (top), peptide crosslinking (middle), and anhydrous muropeptide abundance (bottom) for bifid mutants (*Bbr_0131*, *Bbr_0177*, *Bbr_0431*, *Bbr_1369*, *Bbr_1372*, *Bbr_1373*, and *Bbr_1800*) in MRS (blue), wild type in MRS (black, negative control), and wild type in MRS* (red, positive control). Total peptidoglycan was not correlated with bifid cell abundance (*r*=-0.005, *p*=1). Peptide crosslinking was significantly positively correlated with bifid cell abundance (*r*=0.5, *p*=0.003). The fraction of anhydrous muropeptides, which is inversely proportional to glycan length, was correlated with bifid cell abundance (*r*=0.5, *p*=0.02), but not if data from MRS* was excluded (*r*=0.04, *p*=0.9). *r*: Pearson’s correlation. Bifid cell shape mutants were enriched for fitness defects during colonization and during growth *in vitro*. Plotted is the normalized enrichment score of the 24 bifid mutants during colonization, as measured by fitness scores from the mutant fitness compendium, or during growth *in vitro*, as measured from normalized maximum growth rate (*µ*_max_), maximum OD_600_, and lag time (A) extracted from individual growth curves.

We next measured growth curves and stationary-phase cell shape of the ordered collection in three conditions: MRS, RCM, and MRS supplemented with 300 mM NaCl. By quantifying cell morphologies for every strain in the ordered collection in high throughput^68^, we identified 43 mutants with non-rod-like shapes in at least one growth environment, including bulged poles, bifid shapes, increased cell width, decreased cell length, and “lumpy” (variable cell width) morphologies (**Figure S5**). Of these mutants, 24 exhibited increased frequency of bifid cell shape in at least one growth condition (**Figure 5C**), including genes predicted to be involved in lipid modification (*Bbr_1038*), small molecule transport (*Bbr_1669*), rRNA methylation (*Bbr_1911*), and chromosome segregation (*Bbr_1920*). The breadth of biological processes associated with the bifid phenotype suggest that the factors influencing it are more complex than cell wall synthesis alone. However, since these mutants may have pleomorphic phenotypes that indirectly impact cell wall synthesis, we sought to directly test the impact of a subset of mutations on cell wall architecture.

We profiled the cell wall architecture of seven mutants (*Bbr_0131*, *Bbr_0177*, *Bbr_0431*, *Bbr_1369*, *Bbr_1372*, *Bbr_1373*, and *Bbr_1800*), all of which displayed some degree of bifid cell shape (**Figure S5B**) and shared strong negative phenotypes with cell wall– targeting drugs. We used wild-type cells grown in MRS as a negative control **(Figure 5B)** and wild-type cells grown in MRS*, the sub-optimal formulation that induces aberrant cell shapes (**Figure 5B**), as a positive control. We examined the correlation between bifid cell abundance and three features of cell wall architecture: total peptidoglycan, crosslinking, and the abundance of anhydrous muropeptides, a proxy for peptidoglycan chain length.

The total amount of peptidoglycan varied little between samples, despite the abundance of bifid cells changing by more than an order of magnitude (**Figure 5D**). The fraction of peptidoglycan crosslinking was significantly correlated with bifid cell abundance (*r*=0.5, *p*=0.003; Pearson’s correlation) **(Figure 5D).** The anhydrous fraction, an inverse proxy of peptidoglycan chain length, was correlated with bifid cell abundance (*r*=0.5, *p*=0.02; Pearson’s correlation). However, the correlation coefficient decreased nearly to zero if data from wild-type cells in MRS* were excluded (*r*=0.04, *p*=0.9; Pearson’s correlation). In total, we found no significant connection between branching and decreased cell wall synthesis, either through lower total peptidoglycan, decreased crosslinking, or shorter peptidoglycan chain length.

An opposing hypothesis regarding bifid cell shape is that it is an adaptive response that provides a fitness advantage under stressful conditions^64,69^. To test this hypothesis, we mined our large-scale datasets for an association between bifid mutants and three distinct fitness measurements: growth *in vitro*, stress resistance, and colonization *in vivo*. The bifid mutants were enriched for poor growth *in vitro* in both MRS and RCM (**Figure 5E**), as measured from growth parameters extracted from the individual growth curves (**Methods**). Consistent with this observation, the bifid mutants were significantly enriched for negative phenotypes in 253 *in vitro* conditions while only significantly enriched for positive phenotypes in two: nitrofurantoin and novobiocin (**Table S6**). Finally, we found that the bifid mutants were enriched for poor colonization *in vivo* across all combinations of host and diet (**Figure 5E**).

The breadth of biological functions associated with bifid mutant genes suggests that there are multiple paths to bifid cell formation. While we have shown that the cause of bifid cell shape is more subtle than being a death phenotype associated with cell wall disruption, the pervasive fitness defects associated with bifid cell formation indicate that branched cells are not an adaptive advantage in most environments.

### Parallel pathways of aromatic lactic acid production recycle redox cofactors

Lactic acid metabolites derived from aromatic amino acids are produced in fermented foods^70^ and in the gut microbiome^9,71,72^. These lactic acid derivatives signal to the immune system through binding to receptors like hydroxycarboxylic acid receptor 3 (HCA_3_)^73,74^ and aryl hydrocarbon receptor (AhR)^75,76^. Indole-3-lactic acid signaling through AhR may be particularly potent in the immature gut epithelium of infants^26^ and is correlated with *Bifidobacterium* abundance in infants of distinct geographical regions^9,71,72^.

While the chemical transformation from amino acid to lactic acid likely involves two sequential reactions, the first catalyzed by an amino acid transaminase and the second by a dehydrogenase (**Figure 6A**), the exact pathway and the enzymes involved in Bifidobacteria remain to be determined. In *Bifidobacterium longum*, a single dehydrogenase enzyme, *aldh*, is required for all three aromatic lactic acid metabolites^72^. The physiological role of aromatic lactic acid production in Bifidobacteria also remains to be demonstrated; an *aldh* mutant strain of *B. longum* had no clear growth phenotypes in rich media^72^. Therefore, we sought to use the *Bb* ordered mutant collection to reveal the genetic architecture of aromatic lactic acid production in *Bb* and to determine the physiological role of these metabolites.

**Figure 6:**
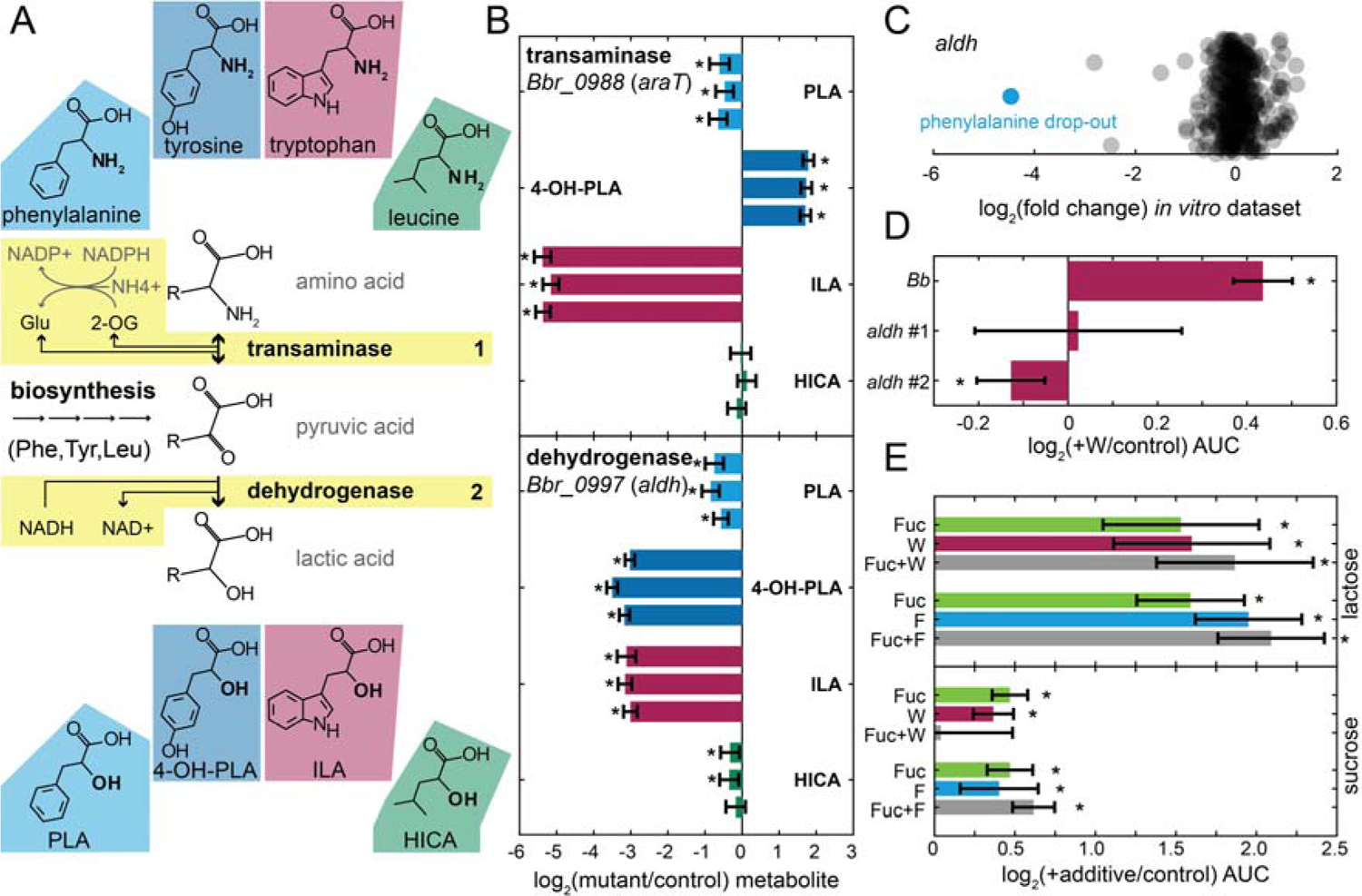
Production of aromatic lactic acids contributes to redox cofactor balance. A) The proposed aromatic lactic acid pathway in *Bb*. Amino acids are converted first to a substituted pyruvic acid by a transaminase enzyme (step 1). Substituted pyruvic acids are converted to substituted lactic acids by a dehydrogenase enzyme (step 2). Phenylalanine is converted into 3-phenyllactic acid (PLA, light blue). Tyrosine is converted into 3-(4-hydroxy)phenyllactic acid (4-OH-PLA, dark blue). Tryptophan is converted into indole-3-lactic acid (ILA, burgundy). Leucine is converted into 2-hydroxyisocaproic acid (HICA, green). The transaminase (step 1) may convert 2-oxoglutarate into glutamate, or an equivalent reaction from keto acid to amino acid. In high ammonium conditions, the action of glutamate dehydrogenase to fix inorganic ammonium would require the conversion of a NADPH to NADP+. The dehydrogenase (step 2) may directly convert NADH to NAD+ as part of the forward reaction. For phenylalanine, tyrosine, and leucine, but not tryptophan, the substituted pyruvic acid is the penultimate metabolite, and the transaminase (step 1 in reverse) is the final enzymatic transformation of biosynthesis. B) An untargeted metabolomic screen of spent media identified a transaminase and a dehydrogenase as contributing to the production of some aromatic lactic acids. Plotted are the log_2_(fold change) values in metabolite levels between the mutant and the pool control (all spent media in the screen pooled together). Data from independent insertion strains (*n*=3) are plotted as individual bars. Error bars represent the 95% CI from technical replicates (1.96×SE, *n*=6). The growth medium was CDM1 with 1% glucose. ILA levels in the spent media of the transaminase *Bbr_0988* (*araT*) mutant were <5% of the pool. AraT was also identified as the biosynthetic enzyme for phenylalanine and tyrosine. The levels of ILA and 4-OH-PLA in the spent media of the dehydrogenase *Bbr_0997* (*aldh*) mutant were <15% of the pool, while the levels of PLA were less impacted. *: *p*<0.05. 95% CI: 95% confidence interval, SE: standard error, *n*: number of replicates. C) In the mutant fitness screen, the strongest fitness defect of *aldh* mutants was during growth in defined media lacking only phenylalanine. Shown are all log_2_(fold-change) values of *aldh* in the *in vitro* fitness compendium. Phenylalanine drop-out is CDM1 with 9 g/L potassium acetate, 1% (w/v) glucose, and all amino acids except phenylalanine. D) Aromatic lactic acid production is the major contributor to the growth boost of 10 mM tryptophan. The area under the curve (AUC) of growth curves was used as a metric of growth. Plotted is the log_2_(AUC ratio) of adding 10 mM tryptophan (+W) to the control medium (defined medium with 0.5% lactose and lacking all amino acids except cysteine and phenylalanine). Error bars represent the 95% CI (1.96×SE, *n*=4). *aldh* mutants, which have curtailed ILA production, do not exhibit the same growth boost as wild type. *: p<0.05 E) Co-metabolism of fucose and aromatic amino acids does not improve growth substantially beyond metabolism of either alone. AUC was used as a metric of growth. Plotted is the log_2_–AUC ratio of adding 0.5% w/v fucose (Fuc), 10 mM tryptophan (W), 10 mM phenylalanine (F), or combinations therein (Fuc+W, Fuc+F) to the control medium; CDM2 with 2% sugar and lacking all amino acids except cysteine. Error bars represent the 95% CI (1.96×SE, *n*=5). The carbohydrate added was either lactose (top) or sucrose (bottom). In lactose media, either fucose or aromatic amino acids provided a significant boost to growth, but co-metabolism was not much greater than from either component alone. In sucrose media, the growth boost from either fucose or aromatic amino acids was lower than in the corresponding lactose medium, and the combination provided a similar boost to either component alone.

To determine the range of metabolites produced by *Bb*, we grew wild-type cultures in a defined complete medium and used an untargeted metabolomic approach to identify metabolites enriched in the spent medium. In addition to 3-phenyllactic acid (PLA) and indole-3-lactic acid (ILA), we detected the lactic acid metabolites derived from tyrosine and leucine, 3-(4-hydroxyphenyl)lactic acid (4-OHPLA) and 2-hydroxyisocaproic acid (HICA), respectively (**Figure 6A, Table S7**). All 4 metabolites have been found previously as products of Bifidobacteria^10,72,77^. Having detected these metabolites, we sought to identify the genes responsible for their production. We assembled a set of insertion mutants in genes annotated with transaminase and dehydrogenase functions, representing both steps of the aromatic lactic acid pathway (**Figure 6A**), analyzed their spent media using untargeted metabolomics, and identified mutants that lost the ability to produce aromatic lactic acids.

We identified two genes with a strong influence on aromatic lactic acid production: one transaminase (*araT*) and one dehydrogenase (*aldh*). Transposon insertions in *araT* severely impacted the production of ILA but had minor effects on PLA levels and increased 4-OHPLA levels (**Figure 6B, top**). We had previously identified *araT* as the transaminase required for the biosynthesis of phenylalanine and tyrosine (**Figure S6**), making it unlikely that the specific effect of *araT* on ILA production reflects specificity of the enzyme. Instead, we propose that either an alternative transaminase or the biosynthetic pathways (**Figure 6A**) provide an alternative source of pyruvic acid derivatives for tyrosine and phenylalanine. The only dehydrogenase tested with a strong influence on aromatic lactic acid production was *aldh*, a homologue of the enzyme identified in *B. longum*^72^. However, in contrast to the phenotype of *aldh* in *B. longum*, we were surprised to find that PLA levels were affected much less by the *aldh* mutant in *Bb* (**Figure 6B**). This result suggests that a second dehydrogenase enzyme in *Bb* plays a redundant role in the production of PLA and was especially interesting given the profile of *aldh* in the fitness compendium.

The strongest fitness effect that we observed for *aldh* was during growth in a chemically defined medium lacking phenylalanine (**Figure 6C**). This observation, combined with our metabolomics results, suggested a model in which multiple genes participate in the aromatic lactic acid pathway, providing partial redundancy. Potentially due to impaired transport of tyrosine (**Figure S2F**), phenylalanine and tryptophan are the major substrates of the pathway, and transformation of either tryptophan or phenylalanine into their lactic acid derivative positively contributes to the growth of *Bb*. Within this model, the sensitivity of *aldh* in phenylalanine dropout media is due to a chemical-genetic interaction (genetic loss of Aldh-dependent transformation of tryptophan and chemical removal of phenylalanine as a substrate) that eliminates flux through the aromatic lactic acid pathway and reduces growth.

We confirmed that the aromatic lactic acid pathway contributes to growth in chemically defined media by examining the growth response of wild-type and *aldh* strains to the addition of tryptophan in the environment. Mutants in *aldh* did not exhibit the same fitness benefits as wild type from the addition of 10 mM tryptophan to a defined medium with low amino acid content (**Figure 6D**). This finding confirmed that the production of aromatic lactic acids can provide a fitness benefit to Bifidobacteria and led us to explore the nature of this fitness benefit further.

Predicted byproducts of the aromatic lactic acid pathway (**Figure 6A**) provide some insight into its physiological benefit. The pathway has the potential to save an NADPH at the transaminase step and to directly regenerate NAD+ at the dehydrogenase step. Thus, production of aromatic lactic acids could contribute to recycling of redox cofactors in Bifidobacteria. This strategy resembles the fucose degradation pathway in Bifidobacteria^78,79^, which is capable of regenerating NADPH and NAD+ but not ATP. Given the overlap in their proposed metabolic byproducts, we hypothesized that studying the co-utilization of fucose and aromatic amino acids could provide insights into the mechanism of the fitness benefit of the lactic acid pathway. We used a low amino acid medium to ensure flux through biosynthetic pathways, draw intermediates away from lower glycolysis, and exacerbate the need for alternative redox balancing reactions. Addition of either fucose, tryptophan, or phenylalanine had a strong positive effect on growth in defined media missing most amino acids (**Figure 6E**). However, the addition of both fucose and an amino had no stronger benefit than either alone (**Figure 6E**). The strength of the positive effect of these compounds depended on the carbon source, with a much stronger effect in 0.5% lactose than 0.5% sucrose, but the lack of a multiplicative effect was consistent in both experiments (**Figure 6E**). These results support a model in which aromatic amino acids are metabolized in parallel pathways to regenerate redox cofactors.

## Discussion

In this work we describe the creation of genome-scale mutant resources for *Bifidobacterium breve* (*Bb*) based on randomly barcoded transposon mutagenesis. We demonstrate the utility of the *Bb* fitness compendium by investigating a broad spectrum of biological phenomena, from metabolism to stress resistance, cell shape, and small molecule production. Many discoveries remain to be made in *Bb* and we expect that the compendium will continue to be a valuable resource for Bifidobacterial biology. Furthermore, the accumulation of new data will only improve the fitness compendium as a resource. To facilitate such efforts, aliquots of the barcoded transposon pool and copies of the ordered mutant collection are available to the scientific community, and we have designed the Fitness Browser (https://fit.genomics.lbl.gov/) to allow for continued access and contribution to this community resource.

Evidence suggests that *Bifidobacterium* sp. are a useful probiotic intervention for multiple early life disorders^5–7^. The production of immunomodulatory aromatic lactic acids by Bifidobacteria^9,71,72^ is proposed to be an important feature of these interventions, but the pathway remains poorly understood. We demonstrated a fitness benefit to *Bb* as well as a physiological role for aromatic lactic acid production in redox cofactor balance. However, the importance of aromatic lactic acid production to *Bb* likely depends on multiple aspects of the external environment. In conditions where redox cofactors are not limited, the contribution of the pathway to growth is minimal (**Figure 6E**). The identity of the co-metabolized carbohydrate also appears to play an important role (**Figure 6E**). While the mechanism behind this effect is unclear, it is important to note that other pathways exist in *Bb* to balance both NAD+ and NADPH. The extent to which carbon source influences flux through alternative pathways and controls the total balance of waste products remains to be determined. It will be interesting to test whether defining environmental conditions according to *Bb* physiology is an effective approach for tuning the amount and relative balance of aromatic lactic acids. Fully elucidating the genetic architecture supporting aromatic lactic acid production in *Bb* will also be useful for understanding the variation observed across *Bifidobacterium* species in terms of the production of these important immunomodulatory small molecules.

New approaches are needed to mine the vast wealth of molecular biology employed across the microbial world. The predictive power of studying mechanisms in model organisms degrades with evolutionary distance and disappears completely when studying novel behaviors not represented by a model. At the same time, the sheer number and diversity of microbial species with important phenotypes makes it virtually impossible to employ the same effort and time that supported the decades of research into models like *E. coli* spread across hundreds of labs. The method that we employed here (**Figure 1A**) represents a highly impactful new approach to bringing mechanistic investigations to the broader microbial world. Focusing on *Bifidobacterium breve* (*Bb*), we showcased the power of a large-scale fitness compendium to accelerate our understanding of non-model organisms. When sequence predictions of gene function were present in *Bb*, the fitness compendium provided experimental confirmation of the predictions (**Figure 1C**), distinguished the roles of highly similar enzymes (**Figure 1D**), identified substrates of enzymes with non-specific annotations (**Figure 1C,2C**), and clarified mis-annotations (**Figure 2C**). When sequence predictions were missing or non-informative, the fitness compendium identified genes that contribute to phenotypes (**Figure 1C,2-6**) and the ordered mutant collection was an invaluable resource for accelerating mechanistic investigations into gene function (**Figure 2-6**). We expect the fitness compendium and ordered mutant collection to be important resources for future research into Bifidobacterial biology and for this study to serve as a template for the rapid characterization of other non-model microbes.

## Supporting information

Dataset S1: The mutant fitness compendium.

Dataset S2: The ordered mutant collection.

Dataset S3: Metabolomics screen of aromatic lactic acid pathway.

Table S1: Plasmids and oligos used in this study.

Table S2: Strains used in this study, beyond experiments performed with the transposon pool and ordered collection.

Table S3: Composition of media used in this study.

Table S4: Muropeptide species detected in UPLC-MS.

Table S5: Strains with aberrant cell shapes.

Table S6: GSEA enrichments for branched cells in the fitness compendium.

Table S7: Metabolites produced by Bb.

## Supplementary Figures

**Figure S1:**
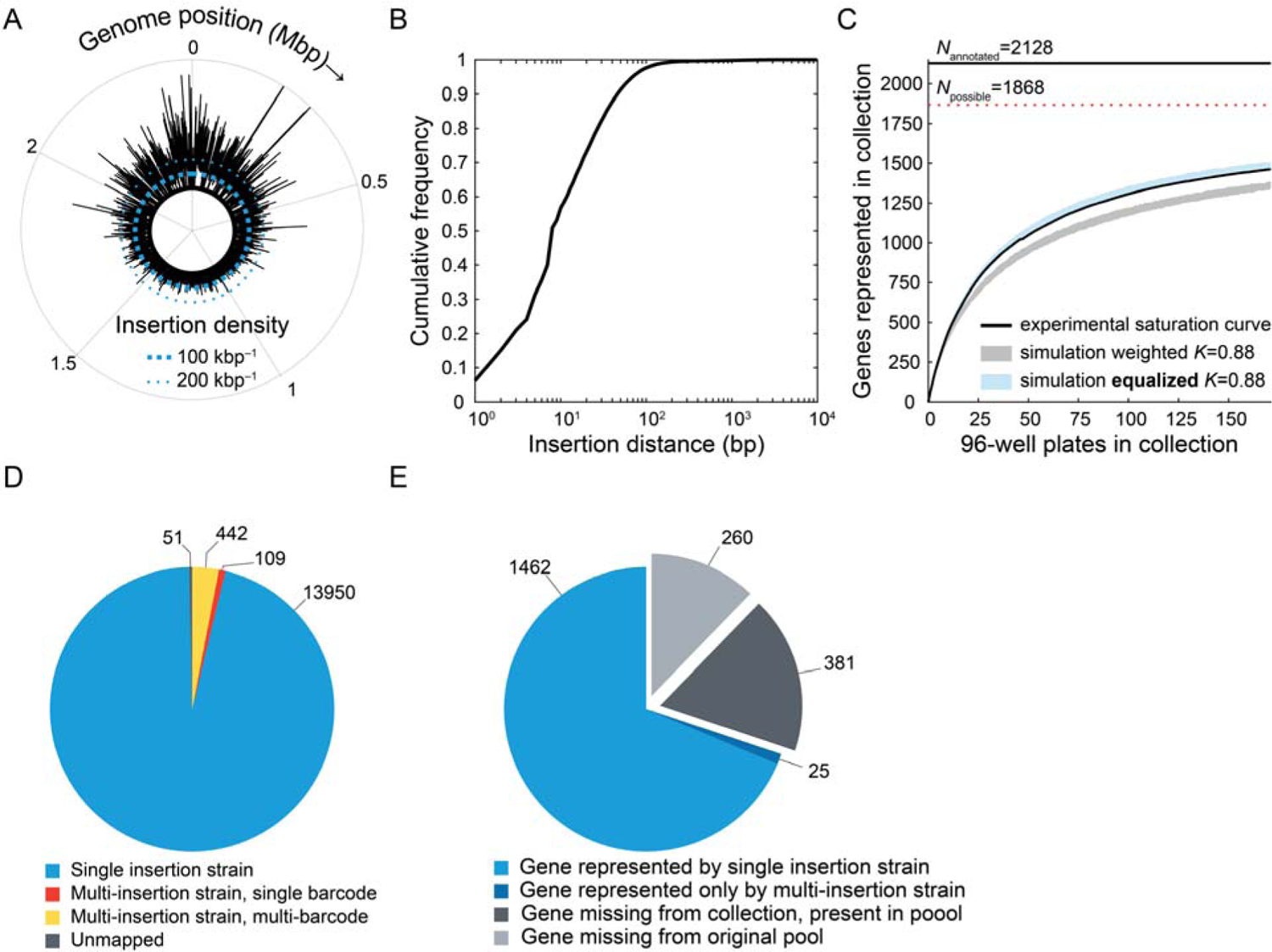
Statistics of transposon collection assembly. A) The density of transposon insertions across the *Bb* genome in the pool. The polar coordinate represents genome position in Mbp, starting at 90° and increasing clockwise from 0 to 2,422,684 bp. Transposon insertion density (kbp^-1^, 250-bp bins) is plotted as function of genome position with markers for 0 kbp^-1^ (thick black line), 100 kbp^-1^ (thick blue dashed line), and 200 kbp^-1^ (thin blue dashed line). Insertion density was higher near the origin (0 Mbp), likely reflecting multi-fork replication at the time of transformation. B) Cumulative frequency distribution of inter-insertion distances. All zero-distance intervals, corresponding to two insertions that co-occur at the same location, were excluded from this analysis. More than half of all inter-insertion intervals are 8 bp or shorter. C) The measured saturation curve of the progenitor collection compared to computational predictions. A Monte-Carlo approach described previously^80^ was used to predict the saturation curve (coverage of the genome as a function of progenitor collection size) using BarSeq data and statistics of the progenitor collection. The assembly efficiency factor *K* (single-insertion strains isolated per well of the progenitor collection) was calculated *ex post facto* to be 0.88. The 95% confidence interval is plotted for simulations in which strain abundance was weighted by control samples from the fitness compendium (weighted, light grey) or artificially set to equal abundances (equalized, light blue). The actual saturation curve is shown as a black line. The equalized simulation better represented the actual saturation curve, likely reflecting the shorter recovery period of the cryo-stock during the sorting experiment as compared to the overnight recovery of the control samples (Methods). A solid black horizontal line represents the number of annotated genes (*N*_annotated_=2128) and the dashed red line marks the hypothetical coverage limit (*N*_possible_=1868), which is the number of genes with at least one insertion in the control BarSeq data. D) Strain-level statistics of insertions in the progenitor collection. Barcodes were binned according to exact co-occurrence in the progenitor collection, and these bins were treated as effective “strains”^80^. Single insertion strains (*n*=13,950) are bins with only one barcode and for which that barcode is associated with only one insertion site. Multi-insertion single barcodes strains (*n*=109) are bins with only one barcode and for which that barcode is associated with multiple insertion sites. Multi-insertion multi-barcode strains (*n*=442) are bins for which multiple barcodes co-occurred in the progenitor collection. Unmapped strains (*n*=51) are barcodes without a corresponding insertion site. Most strains in the *Bb* progenitor collection are single-insertion strains. Gene-level statistics of coverage in the progenitor collection. Annotated genes (*n*=2128) are classified according to representation in the progenitor collection according to the classification in (D). These include genes represented by at least one single-insertion strains (light blue, *n*=1462), genes represented only by strains with multiple insertions (dark blue, *n*=25), genes missing from the collection but present in the pool (grey, *n*=381), and genes missing from the original pool (black, *n*=260).

**Figure S2:**
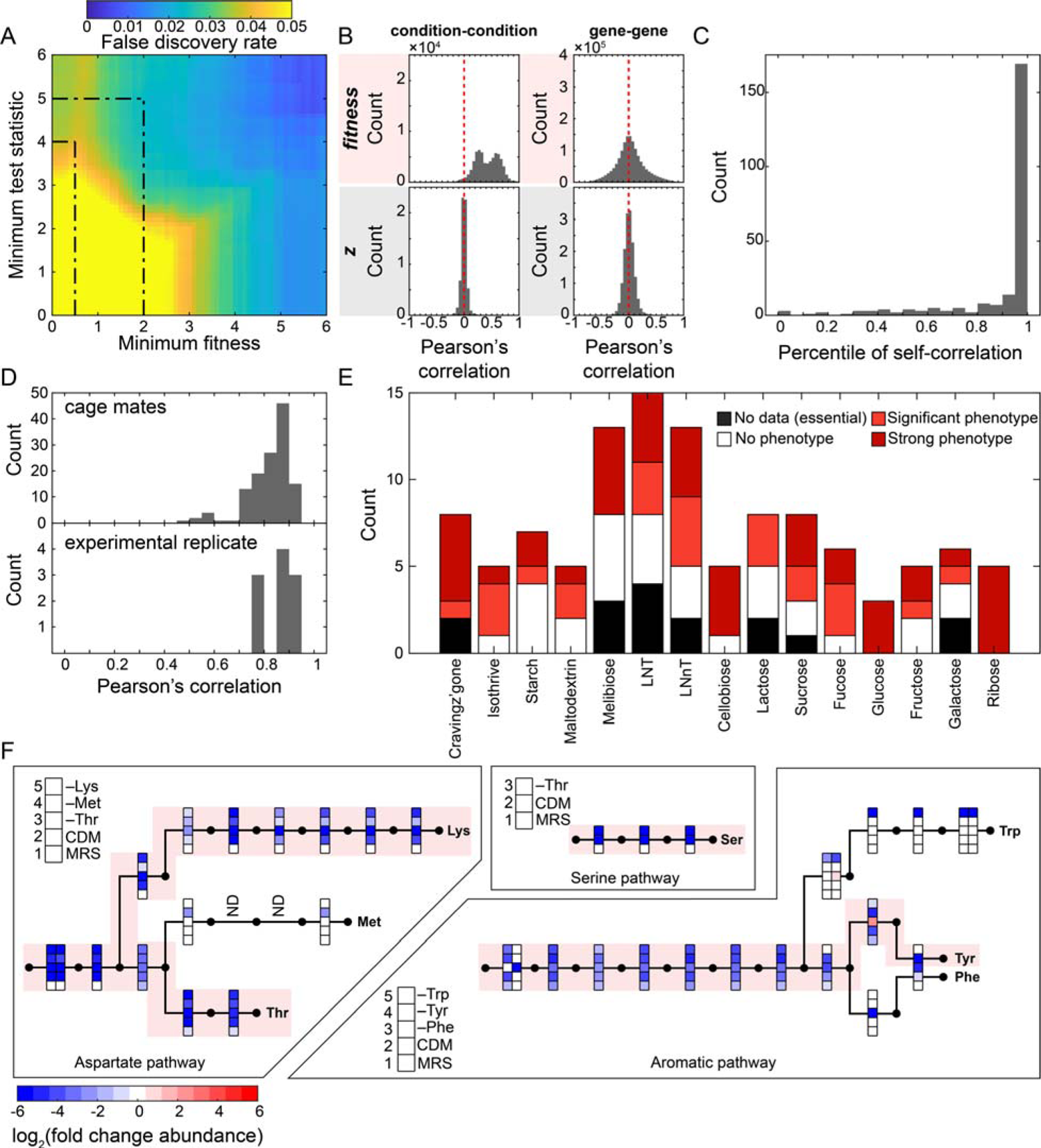
The mutant fitness compendium is reproducible and rich in biological information. A) An estimate of the false discovery rate (FDR) as a function of *fitness* and *t*. A non-parametric estimate of the FDR for fitness phenotypes was calculated from control and experimental (non-control) samples in the mutant fitness compendium (**Methods**). The cutoffs defined previously for significant (|*fitness*|>0.5 and |*t*|>4) and strong (|*fitness*|>2 and |*t*|>5) phenotypes^19^ have FDR<0.05. Batch normalization of fitness scores reduces the baseline condition-condition and gene-gene correlations. Batches are defined as the samples collected at the same time, prepped in the same library, and sequenced on the same run. Fitness phenotypes (top row, red) accurately represent strain fitness but suffer from batch effects that lead to high intra-batch correlations between conditions (top left) and a more variable distribution of gene-gene correlations (top right). Batch normalization (bottom row, grey) reduces the intra-batch correlation (bottom left) and variability of gene-gene correlations (bottom right) (**Methods**); the resulting *z*-score is a chemical interaction score instead of a direct measurement of fitness^81^. Fitness phenotypes were used for most analyses, but *z*-scores were used for hierarchical clustering of genes and other correlation-based measures. C) Measurements from the *in vitro* dataset are highly reproducible between replicates. Correlations were calculated both between replicate measurements in a batch and between a sample and every non-replicate measurement within the batch. If >2 replicates were measured, the most correlated pair was saved. The data used for these correlations were the *z*-scores. Plotted are the percentile of replicate correlations among replicate and non-replicate measurement. A percentile of 0.95 indicates that a replicate correlation is more correlated than 95% of the non-replicate comparisons. Most replicate correlations are above the 90^th^ percentile. D) Replicate measurements from the *in vivo* dataset are highly correlated. Top: the distribution of Pearson’s correlations between cage mates (same experiment, diet, and day; mice and chicks are both considered). These measurements were subsequently averaged in the mutant fitness compendium. Bottom: the distribution of Pearson’s correlations for comparisons between the first (male mice) and second (female mice) mouse colonization experiments, matched by day and diet. E) The fitness compendium informed models of carbohydrate utilization pathways in *Bb*. Genes for small molecule transport and CAZymes were assembled into pathways based on sequence homology and fitness phenotypes. Most pathways include initial import and enzymatic steps to a phosphorylated precursor fed into glycolysis: glucose-6-phosphate or fructose-6-phosphate. The fucose pathway includes full transformation to the waste product 1,2-propanediol. Genes are grouped in pathways and colored according to fitness phenotypes during growth on specific carbohydrates. Color summarizes genes with strong fitness phenotypes on the carbohydrate (dark red), with significant fitness phenotypes on the carbohydrate (light red), predicted by sequence homology and with no significant fitness phenotypes (white), and predicted by sequence homology and with no data in the fitness compendium (black). F) Fitness phenotypes associated with serine, tyrosine, threonine, and lysine biosynthesis pathways indicate uptake of the free amino acids is limited in *Bb*. Genes in the aspartate pathway (left), aromatic amino acid pathway (right), and serine pathway (top) are arranged according to position in the pathway. Pathway intermediates are represented as filled circles and enzymatic steps are represented as arrows connecting the circles. Relevant fitness phenotypes are plotted as a heatmap for each gene at the position of its enzyme within the pathway. The bottom two fitness phenotypes are MRS and CDM1 (with all amino acids supplemented) for each pathway, and the remaining data are drop-out experiments of the relevant amino acids to each pathway. Mutants within a pathway consistently unable to grow on CDM1 indicate that uptake of the associated amino acid is insufficient to support growth. Despite sharing the initial steps of the pathway, lysine and threonine can be distinguished from methionine (aspartate pathway, left) and tyrosine can be distinguished from phenylalanine and tryptophan (aromatic amino acid pathway, right).

**Figure S3:**
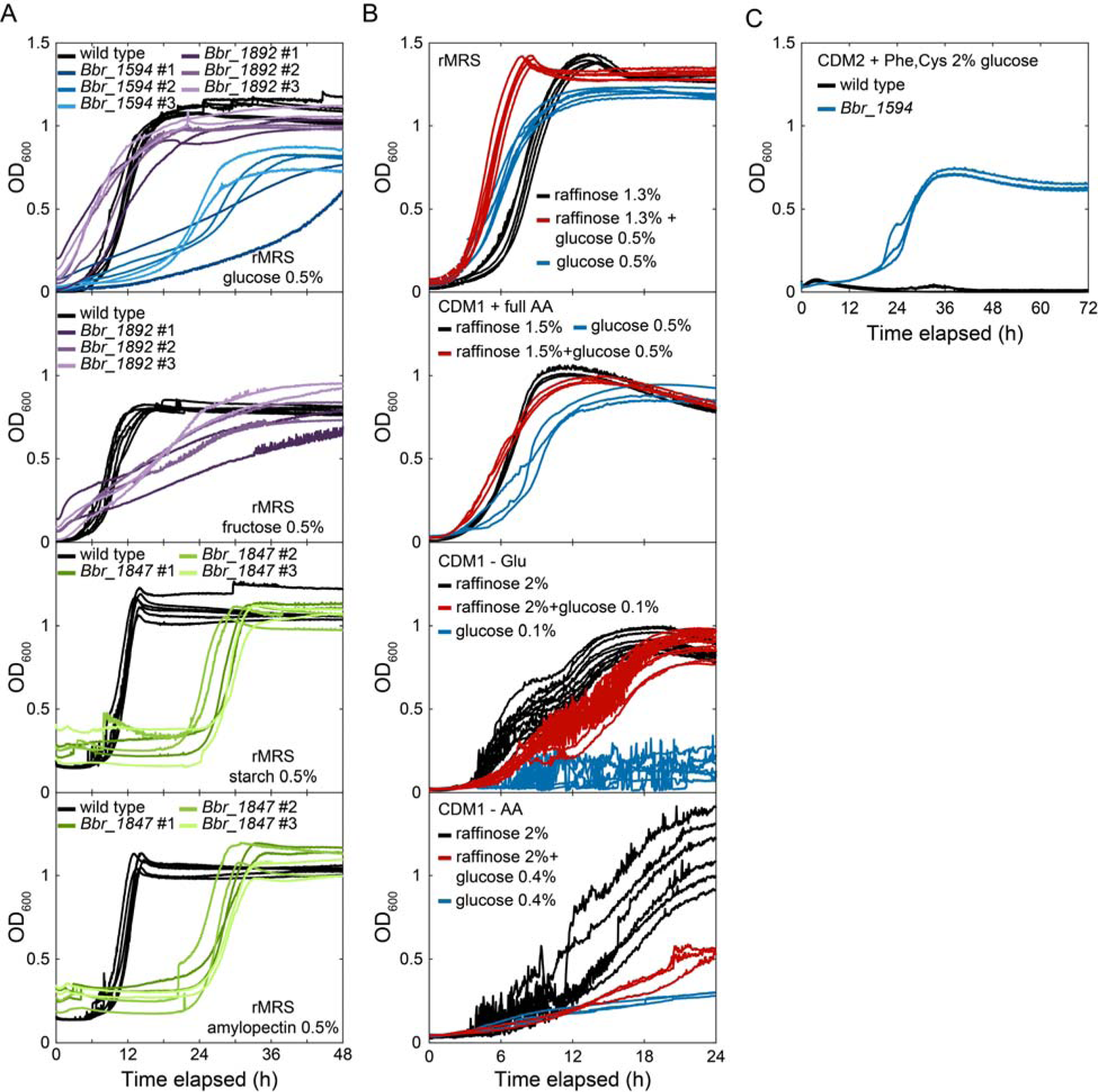
Growth behavior of sugar transport mutants *in vitro*. A) Mutant growth curves confirm predicted substrates of sugar transporters. Shown are growth curves for individual cultures, colored by genetic background. Top: wild type (black, *n*=6), insertion strains in *Bbr_1594* (shades of blue, *n*=2), and insertion strains in *Bbr_1892* (shades of purple, *n*=2) were grown in rMRS with 0.5% (w/v) glucose. The growth defects of *Bbr_1594* insertion strains are greater than those of *Bbr_1892* strains, supporting the conclusion that *Bbr_1594* encodes the main glucose transporter in *Bb*. Second from top: wild type (black, *n*=6) and insertion strains in *Bbr_1892* (shades of purple, *n*=2) were grown in rMRS with 0.5% (w/v) fructose. The growth defects of *Bbr_1892* in fructose are greater than those in glucose, supporting the conclusion that fructose, and not glucose, is an important substrate of the transporter encoded by *Bbr_1892*. Second from bottom: wild type (black, *n*=6) and insertion strains in *Bbr_1847* (shades of green, *n*=2) were grown in rMRS with 0.5% (w/v) starch. The insertion strains exhibited a growth phenotype, supporting the conclusion that the transporter encoded by *Bbr_1847* imports starch. Bottom: wild type (black, *n*=6) and insertion strains in *Bbr_1847* (shades of green, *n*=2) were grown in rMRS with 0.5% (w/v) amylopectin. The insertion strains exhibited a growth phenotype, supporting the conclusion that the transporter encoded by *Bbr_1847* imports amylopectin. B) The presence of glucose in the environment has increasingly negative effects in media with decreasing amino acid content. Shown are growth curves for individual wild-type cultures in media supplemented with raffinose (black), glucose (blue), and a combination of raffinose and glucose (red). Top: in rMRS, a rich medium with free amino acids and peptides, glucose supports faster growth than raffinose and a combination of sugars grows better than either sugar alone (*n*=6, each condition). Second from top: in CDM1 with all 20 free amino acids, raffinose supports faster growth than glucose, and a combination of the two sugars performs similarly to raffinose alone (*n*=3, each condition). Second from bottom: in CDM1 without glutamate (-E), glucose supports very little growth alone and the combination of glucose and raffinose performs worse than raffinose alone (*n*=6, 12, 12 for glucose, raffinose, and glucose+raffinose, respectively). Bottom: in CDM1 lacking all amino acids except cysteine (-AA), glucose supports very little growth and the combination of glucose with raffinose also grows very little (*n*=3, 6, 3 for glucose, raffinose, and glucose+raffinose, respectively). C) The lack of growth due to glucose in nutrient-poor media is likely not due to carbon catabolite repression, because a mutation in the glucose transporter *Bbr_1594* rescues growth in media with glucose as the only carbon source. Shown are individual growth curves for wild type (black) and a *Bbr_1594* insertion strain (blue) in CDM2 with only cysteine and phenylalanine as amino acids and 2% (w/v) glucose (*n*=3). Growth of *Bbr_1594* under this condition suggests that secondary transporter exists for glucose and that glucose transported by *Bbr_1594* is directly inhibiting growth of *Bb*.

**Figure S4:**
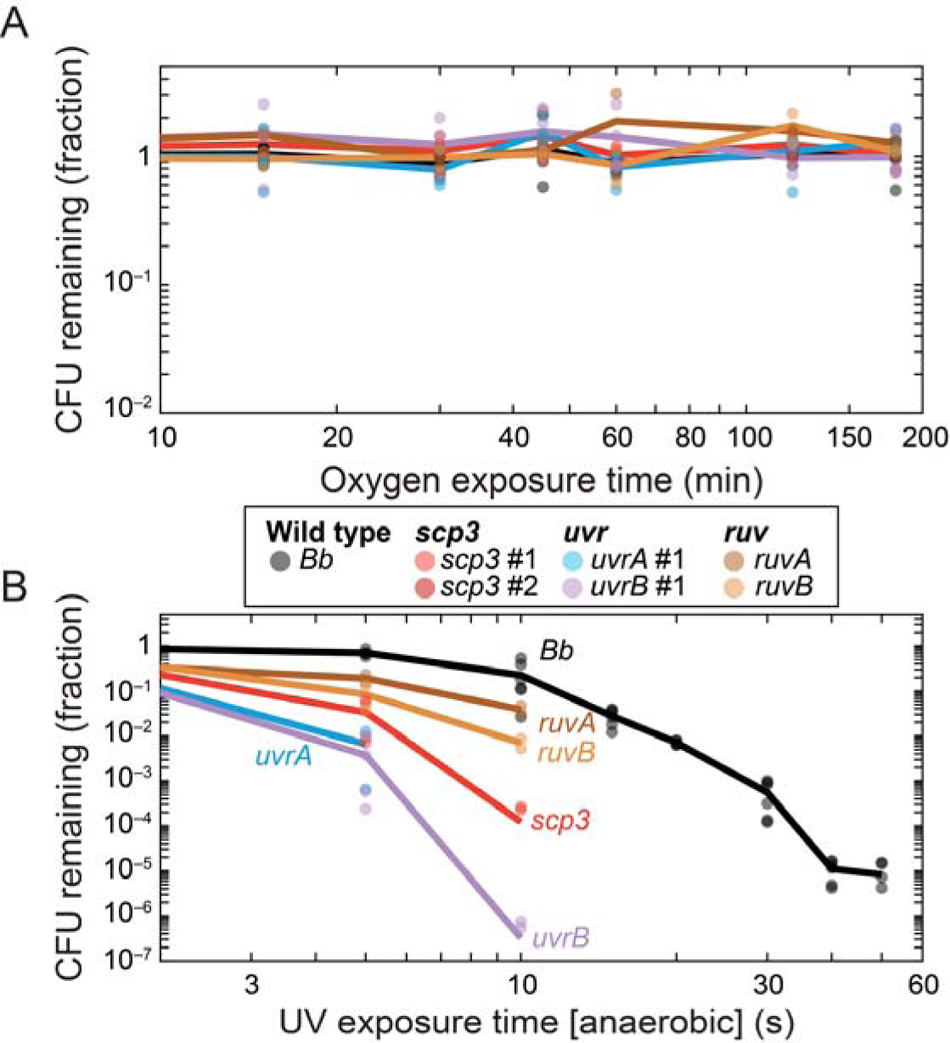
*scp3*, *uvr*, and *ruv* strains are not sensitized to oxygen. A) The fraction of colony forming units (CFU) remaining as a function of oxygen exposure time. CFU did not decrease substantially after 3 h of aerobic incubation at 37 °C for any of the strains. B) The fraction of CFU remaining as a function UV exposure time in an anaerobic chamber (**Methods**). The results are qualitatively similar to data in Figure 4C, which were collected in aerobic conditions. These data support the conclusion that *scp3* is specifically sensitive to UV, rather than a combination of UV and oxygen.

**Figure S5:**
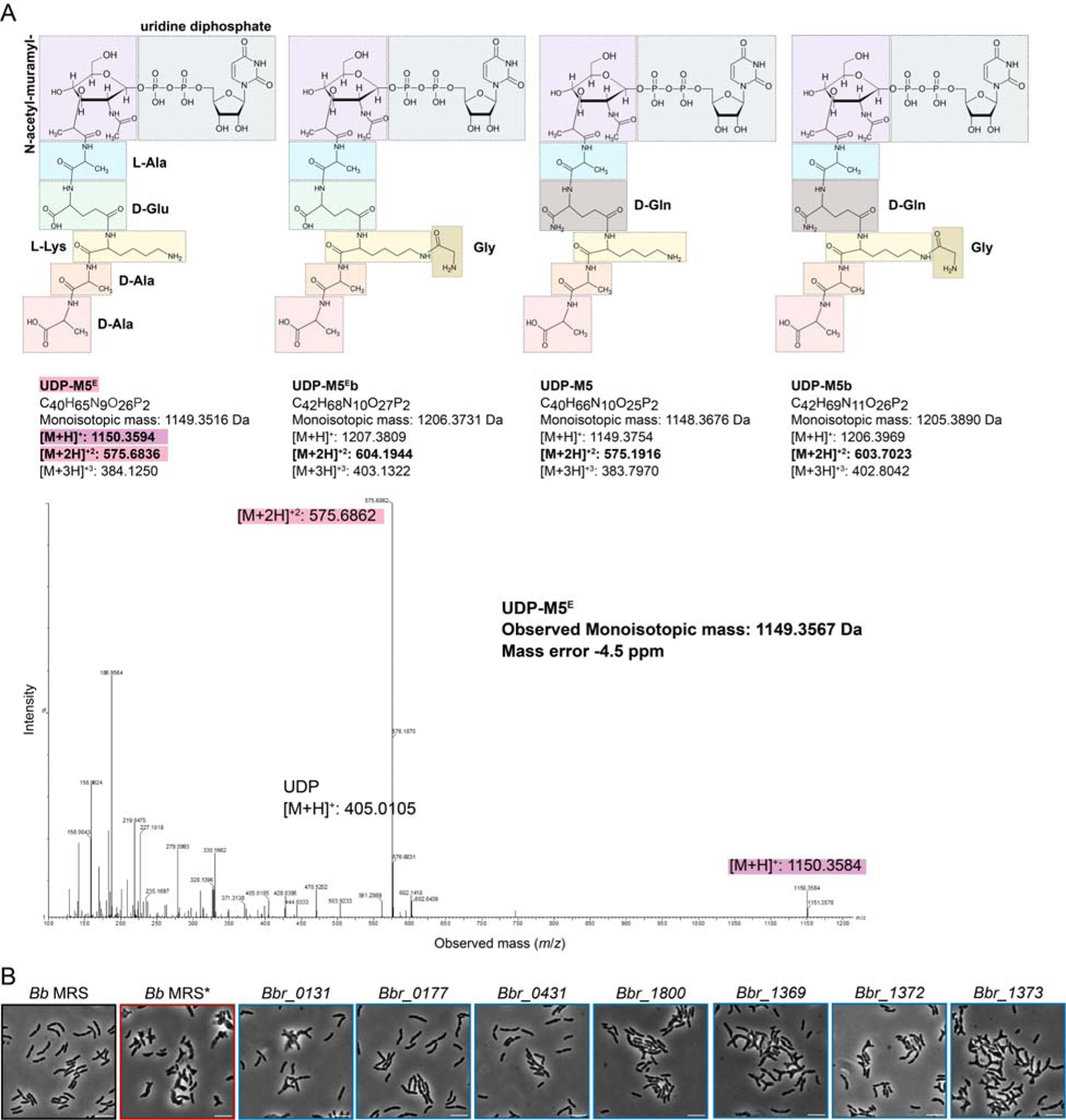
Cell wall features and cell shape defects of *Bb*. A) *m*/*z* values of the soluble precursor to the M5 monomer are consistent with UDP-M5^E^, a stem-peptide lacking both the glycine crossbridge and the amidation of D-isoglutamate. Top: hypothetical muropeptide precursors vary at both the amidation of D-glutamate at position 2 (D-Glu | D-Gln) and the addition of a glycine crossbridge at position 3. Bottom: the MS/MS profile of untargeted MS in the cytosolic fraction of *Bb* lysate. *m*/*z* peaks associated with the [M+H]^+^ and [M+2H]^+^ ions are consistent with UDP-M5^E^. Observation of UDP-M5^E^ indicates that these two modifications are added after synthesis of the lipid II precursor in *Bb*. B) Representative images of single cells of the seven mutants (blue outline) analyzed for cell shape and cell wall features, along with negative (*Bb* MRS, black outline) and positive (*Bb* MRS*, red outline) controls. Images were taken of cells in stationary phase after 17 h of growth. The scale bar is 5 µm.

**Figure S6:**
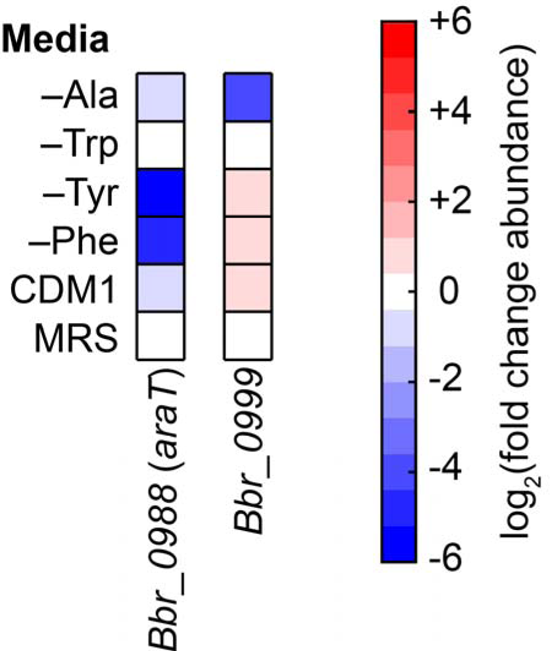
*araT* is involved in aromatic amino acid biosynthesis. Heatmaps represent the fitness values of two genes annotated as amino acid transaminases, *Bbr_0988* (*araT*) and *Bbr_0999*. Fitness phenotypes in amino acid drop-out conditions connect each gene to their respective biosynthetic pathways. *Bbr_0988* (*araT*) was connected to the synthesis of tyrosine and phenylalanine and to indole lactic acid production (Figure 6B). *Bbr_0999* was previously postulated to play a role in the aromatic lactic acid pathway due to its proximity to *Bbr_0997*(*aldh*) in *Bifidobacterium longum*^72^. *Bbr_0999* was connected to alanine biosynthesis but had no phenotypes involving aromatic lactic acid production.

## Supplementary Datasets

**Dataset S1: The mutant fitness compendium.** Data covers the genome-wide fitness phenotypes, test statistics, and *z*-scores of *Bb*. Genes are in rows, referenced by their locus ID. Experiments are in columns, referenced by the sequencing run (set) and index number (IT) (e.g., set3IT004). Data that was averaged across multiple experiments has a column name corresponding to the perturbation. Detailed experimental information is provided for each data set as a metadata table.

**Dataset S2: The ordered mutant collection.** Included is strain location in the condensed collection (FinalPosition), strain location in the progenitor collection (SourcePosition), the molecular barcode of the transposon insertion (Barcode), the scaffold (Scaffold), the position in the genome (Position), the orientation of the transposon (Orientation), the gene disrupted (Gene), notes on potentially overlapping barcodes (Overlap), and the number of times the barcode occurs in the progenitor collection (PickCount).

**Dataset S3: Metabolomics screen of aromatic lactic acid pathway.** Quantification of 7 metabolites (columns) across multiple mutant strains, with six replicate measurements per strain (rows). The alignment ID (Alignment ID), average retention time in minutes (Average Rt(min)), average *m*/*z* (Average Mz), metabolite name (Metabolite name), and INCHIKEY are recorded for each measured metabolite. The internal standards D3-glutamic acid, D5-glutamine, and D5-hippuric acid are included but were not analyzed.

## Supplementary Tables

**Table S1: Plasmids and oligos used in this study.** For oligos, the description, sequence, purpose for this study (purpose), and molecular target (target) are provided. For plasmids, the plasmid name (plasmid), antibiotic resistance, and the source of the plasmid (source) are provided.

**Table S2: Strains used in this study, beyond experiments performed with the transposon pool and ordered collection.** The strain name (Strain), ID of parental strain (Derived from ID), gene name #1 (UID1), gene name #2 (UID2), and antibiotic resistance cassette associated with the mutation (resistance cassette) are provided. If the strain is a transposon insertion from the collection, barcode sequence, insertion position, insertion orientation, and the strain designation are provided. We use the location in the progenitor collection as a strain designation. Also provided are the names of plasmids in the strain, the antibiotic resistances, the source of the strain, and the associated figure name (associated data) in which the strain was used.

**Table S3: Composition of media used in this study.** The composition of the two chemically defined media used in this work. The compound name (Compound), manufacturer, and catalog number (Cat. No.) are provided. The concentration is provided in g/L except for polysorbate 80, which is recorded in mL/L (denoted by asterisk).

**Table S4: Muropeptide species detected in UPLC-MS.** The chemical composition of muropeptide species detected in *Bb*. The retention time in minutes (RT (min)), short name (Name), and chemical structure (Structure) are provided, along with the theoretical (Theoretical) and observed (Observed) neutral mass in Daltons and the difference between theoretical and observed masses (Difference).

**Table S5: Strains with aberrant cell shapes.** Cell shape classification of the ordered collection. The position in the ordered collection (FinalPosition) and ID of disrupted gene (Gene) are provided, along with the manually classified shape phenotype and severity in MRS, RMC and MRS+salt. Only strains with aberrant shapes are included.

**Table S6: GSEA enrichments for branched cells in the fitness compendium.** Gene Set Enrichment Analysis (GSEA), testing the 24 mutants identified as bifid for enrichment for positive or negative fitness phenotypes in the *in vitro* growth conditions. Provided are the name of the bifid cell group (NAME: BRANCH_ID), the growth condition being tested for enrichment (CONDITION), the direction of the enrichment (VALUE: POSITIVE OR NEGATIVE), the enrichment score (ES) and normalized enrichment score (NES), the nominal *p*-value (NOMP_val), and false discovery rate (FDR). Only conditions with FDR<0.05 are listed. The majority of enrichments of the bifid mutant group are negative.

**Table S7: Metabolites produced by *Bb*.** An annotated set of compounds produced by *Bb*. Data are provided for a chemically defined medium (CDM1, negative control) and the spent medium of *Bb* grown in CDM1. The retention time in minutes (Rt), *m*/*z* value (Mz), and compound name (Name) are provided. Compounds are grouped according to type (e.g. acetylated amino acids and gamma-glutamylated amino acids).

## Materials and Methods

### Oligos, strains, and media

The plasmids and oligos used in this study are presented in **Table S1**. The strains used in this study are presented in **Table S2.** The composition of base media used in this study are presented in **Table S3**. The chemically defined media (CDM1 and CDM2) and reconstituted MRS (rMRS) formulations are given without a carbon source (**Table S3**); the carbon source used in a particular experiment is explicitly stated.

Transformation broth (TB) and reconstituted MRS (rMRS) are modified versions of MRS^82^ made from individual components (**Table S3**). TB includes supplements previously incorporated into media used for transformation^30^. All components of TB and rMRS were added before sterilizing except the carbon source, L-cysteine, manganese sulfate, and magnesium sulfate. Media were brought to a pH of 6.8 before sterilization and pre-reduced in an anaerobic chamber for at least 48 h, after which the final components were added as pre-reduced filter-sterilized stocks immediately before use (Corning, 430320). The TB was sterilized using an autoclave set at 121 °C for 30 min and the rMRS was sterilized using a polyethersulfone (PES) filter membrane with 0.2-µm pore size (Thermo Scientific, 595-4520).

Unless specified otherwise, the commercial formulation of MRS was Difco brand (BD, 288110). The alternative commercial MRS formulation associated with aberrant cell shape (MRS*) in *Bifidobacterium breve* UCC2003 (Figure 5) was Oxoid brand (Thermo Fisher, CM0359). The recovery broth (RB) used during transformation was commercial MRS supplemented with 3 mM L-cysteine (Alfa Aesar, A10389), 150 µM iron(II) sulfate heptahydrate (Sigma-Aldrich, F7002-250G), and 100 µM calcium chloride dihydrate (Fisher Scientific, C79-500). The selective plates used during transformation were commercial MRS with 1.5% (w/v) agar (BD, 214030), 3 mM L-cysteine, 150 µM iron(II) sulfate heptahydrate, 100 µM calcium chloride dihydrate, and 10 µg/mL tetracycline hydrochloride (MP Biomedicals, 103011). The normalization broth (NB) was commercial MRS with 3 mM L-cysteine, 150 µM iron(II) sulfate heptahydrate, 100 µM calcium chloride dihydrate, and 10 µg/mL tetracycline hydrochloride. For RB, selective plates, and NB, the extra components were added as pre-sterilized stocks to the sterilized media immediately before use. RB and NB were sterilized with 0.2-µm PES filters. Selective plate agar media were sterilized using an autoclave set at 121 °C for 30 min.

The wash buffer (WB) was adopted from previous transformation protocols^30^. Briefly, WB is 500 mM sucrose (EMD Millipore, 8550-5KG) and 1 mM citrate (Research Products International, C35025-500.0) brought to a pH of 5.8 before autoclaving at 121°C for 30 min and pre-reducing in an anaerobic chamber for at least 48 h.

The chemically defined media (CDM1, CDM2) (**Table S3**) used in this study were adapted from previous formulations^83,84^. All components were added before sterilization except for the carbon source, L-cysteine, magnesium sulfate, calcium chloride, iron sulfate, zinc sulfate, and sodium bicarbonate. The media were brought to a pH of 7.0, filter sterilized using a cellulose acetate (CA) membrane with a 0.2-µm pore size (Corning, 430320), and pre-reduced in an anaerobic chamber for at least 48 h. The final components were added as pre-reduced filter-sterilized stocks immediately before use. In experiments tracking mutant fitness in defined media with individual components left out, each component of the defined media was independently prepared as a stock solution and filter sterilized using cellulose acetate 0.22-µm filters. The media were then assembled from individual sterile components in the anaerobic chamber.

### Transposome assembly

The *tetW* resistance cassette was PCR amplified from the plasmid pMOD2-tetW-LacI(fos)term^30^. The 5’ end of the reverse primer contained a random barcode with two nucleotide changes in universal primer binding site 2 (U2) that eliminated two restriction sites recognized by the *B. breve* UCC2003 restriction-modification systems^85^. The resistance cassette was amplified using an initial denaturation step at 98 °C for 30 s, 22 cycles of [a denaturation step at 98 °C for 10 s, an annealing step at 56 °C for 20 s, and an extension step at 72 °C for 2 min], and a final extension step at 72 °C for 10 min. The annealing temperature and number of cycles were determined empirically. Template DNA at a concentration of 0.5 ng/µL was amplified using Phusion (NEB, M0530S) in High Fidelity Buffer supplemented with 3% (v/v) DMSO. DMSO was empirically found to be necessary for amplification.

PCR product from multiple reactions was accumulated and gel extracted from a 0.8% (w/v) TAE agarose gel using a Nucleospin Gel and PCR clean-up kit (Macherey-Nagel, 740609.250). Gels were cut without exposing the DNA to UV radiation damage or stain, according to the manufacturer’s instructions. After elution from the purification columns, transposon DNA was ethanol precipitated and resuspended in TE (10 mM Tris, 1 mM EDTA), pH 7.5 at a concentration of 450 ng/uL.

Transposomes were assembled using EZ-Tn*5* according to the manufacturer’s instructions (Illumina, NP92110), except that the starting concentration of transposon DNA was 450 ng/µL instead of 100 ng/µL. The transposome assembly reaction was performed two days before the electroporation and stored at −20 °C until use.

### Generation of the pooled transposon insertion library

*B. breve* UCC2003 (*Bb*) was struck for single colonies from a glycerol stock on MRS agar plates. After two days, a single colony was used to inoculate 100 mL of MRS broth supplemented with 3 mM L-cysteine and the culture was grown overnight (∼13 h) at 37°C. The overnight culture was used to inoculate 250 mL of MRS supplemented with 3 mM cysteine at an initial OD_600_ of 0.4. This culture was grown at 38 °C for 5.5 h to an OD_600_ of 4.9 (mid-stationary phase). The stationary phase culture was used to inoculate 1 L of TB at an initial OD_600_ of 0.4. The TB culture was grown at 38 °C for ∼3 h to an OD_600_ of 1.9. Cells were chilled on ice, centrifuged (Avanti JXN-26 Centrifuge, JLA-10.500 rotor) at 4,500 *g* and 4 °C for 15 min, and washed twice with 300 mL of ice-cold WB. During every pelleting step, centrifuge bottles were closed tightly, and the lid was sealed with Parafilm before removing the centrifuge bottle from the anaerobic chamber. After the second wash, competent cells were resuspended at a calculated OD_600_ of 150 in ice-cold WB.

Five milliliters of competent cells were thoroughly mixed with 100 µL of transposome reaction. Next, 400-µL aliquots of the mixture were aliquoted in pre-chilled, 0.2-cm gap electroporation cuvettes (Fisher brand, FB102) and electroporated using an Electro Cell Manipulator and Safety Stand (BTX Harvard Apparatus, ECM630 and 630B, respectively). To maintain an anaerobic environment during the electroporations, high voltage extension wires (custom made) were fed into the anaerobic chamber to connect the voltage generator outside to the safety stand and competent cells inside. Cells were electroporated using 10 kV/cm, 350 Ω, and 25 µF.

Electroporated cells were immediately recovered in RB prewarmed to 37 °C. After rescue, the entire 5 mL of competent cells were resuspended in a single flask in 50 mL of RB and incubated at 37 °C for 1 h. The RB culture was plated on selective plates (200 µL of culture per plate). The selective plates were incubated for ∼36 hours at 37°C.

After colonies were visible on the selective plates, the plates were scraped and resuspended in 1.4 L of NB pre-warmed to 37 °C. This broth was incubated at 37 °C with stirring to break up cell clumps for 1 h. Six hundred milliliters of pre-warmed 50% (v/v) glycerol were then added to the culture to make 2 L of a 15% (v/v) glycerol stock. The final OD_600_ of this glycerol stock was 12.6. One liter of this culture was aliquoted and sealed in crimp vials, which were flash frozen using liquid nitrogen and stored at −80°C. The second liter of this culture was pelleted, flash frozen, and stored at −20 °C for DNA extraction and transposon insertion mapping.

### Plasmid transformation of *Bb* strains

The protocol for transformation of plasmids into *Bb* was similar to the protocol used for the transposon insertion pool, with a simplified set of media. Cells were grown in commercial MRS, transferred to TB for isolation of competent cells, electroporated in WB, recovered in commercial MRS, plated on commercial MRS with antibiotics (depending on the plasmid), and colonies were inoculated in commercial MRS with selective antibiotics for cryo-stocks. For plasmids based on pNZ44, selection was with 5 µg/mL chloramphenicol.

### DNA isolation

Genomic DNA was isolated from *Bb* cultures using a DNeasy Blood and Tissue Kit according to the manufacturer’s instructions for Gram-positive bacteria (Qiagen, 69506 and 69581). Genomic DNA was isolated from fecal matter and gastrointestinal sections using the DNeasy HTP 96 PowerSoil Kit (Qiagen, 12955-4). All DNA was recovered in ultrapure water without additives and stored at −20 °C.

### RBTn-seq

We used a previously described protocol^21^ with minor modifications for the RBTn-seq step that associated barcodes with transposon insertion sites in the *Bb* genome.

Briefly, genomic DNA isolated from the transposon pool was sheared to ∼300 bp using a Covaris S2 Ultrasonicator (Covaris), custom adapters (**Table S1**) were ligated using a NEBNext DNA Library Prep Master Mix Set for Illumina (New England Biolabs, E6040S), and the transposon-genome junction was amplified with primers (**Table S1**) that incorporated Illumina adaptor sequences to create the final RBTn-seq libraries.

Multiple independent libraries were made for each sample and combined before sequencing. Sequencing was performed using either a MiSeq (v3 reagents) or a HiSeq4000. Libraries were prepped from both the full transposon pool and from the pooled cultures of the progenitor collection because sequencing the progenitor collection has been shown to be more effective at identifying multiple-insertion strains^80^.

### Bar-seq

We used a previously described protocol^21^ with minor modifications to amplify barcodes for barcode sequencing (Bar-seq). Briefly, genomic DNA was used in a single PCR to amplify the barcode sequences and incorporate Illumina adaptor sequences. The oligos used in this work were modified from previous studies^18^ to eliminate a restriction site in U2 found in *Bb*^85^. Bar-seq oligos included an indexing sequence for multiplexing, and we typically pooled 96 sequencing libraries on a Next-seq (Illumina) for sequencing.

### Fitness measurements for *in vitro* perturbations

Experiments to study carbon source utilization used rMRS (**Table S3**). Experiments tracking the omission of individual components of chemically defined media (drop-out experiments) used CDM1 (**Table S3**) with 10 g/L glucose and 0.9 g/L potassium acetate. Experiments involving growth in the spent medium of another strain used CDM1 (**Table S3**) without MOPS buffer and supplemented with a mix of sugars: 20 g/L glucose, 4 g/L fructose, 4 g/L cellobiose, 4 g/L maltose, and 4 g/L lactose. Our approach for measuring fitness phenotypes *in vitro* was largely based on previous efforts^18–20^. Briefly, an entire cryo-stock was thawed, inoculated into 50 mL of commercial MRS with 10 µg/mL tetracycline, and allowed to recover anaerobically overnight at 37 °C.

In most experiments in the fitness compendium, two solutions were prepared at double concentration and then combined for the final growth condition. The first solution was the perturbation (drugs or carbon sources in water, chemically defined media missing a component) and the second was a culture of the transposon pool in the base medium. These “2X” experiments were assembled as follows. First, for antibiotic treatment and carbon source utilization experiments, the added compound was prepared as a double-concentration solution in water. The overnight culture of the transposon insertion pool was then diluted to twice the inoculation density (OD_600_=0.04) in either double-strength commercial MRS (antibiotics) or double-strength rMRS without a carbohydrate (carbon source). Second, for nutrient-dropout experiments, a mixture of every medium component except one was assembled in a 2-mL 96-deepwell plate (Greiner, 780270) at double concentration. The overnight culture was pelleted and washed three times in a double-concentration salt solution composed of medium components not involved in the dropout experiment, then diluted to an OD_600_ twice the inoculation density (OD_600_=0.04) in the same salt solution. One milliliter of the solution containing the perturbation and 1 mL of the initial culture were combined in a 2-mL 96-deepwell plate (Greiner, 780270).

In experiments involving growth in the spent medium of another strain, double strength solutions were not prepared. Instead, a culture was grown in 2 mL of the chemically defined medium in a 2-mL 96-deepwell plate for 48 h. The culture was then pelleted, and the supernatant was removed, stored in a 2-mL 96-deepwell plate, filter sterilized in a 96-well filter plate (AcroPrep Advance 96-well 0.2-μm GHP 636 membrane filter plate, PALL), and stored at −20 °C for future use. On the day of the experiment, the overnight culture was pelleted and washed three times in CDM1b, then resuspended at double density (OD_600_=0.04) in fresh CDM1b. One milliliter of the solution containing the perturbation and 1 mL of the initial culture were combined in a 2-mL 96-deepwell plate (Greiner, 780270).

After assembly of all growth conditions, the 96-deepwell plate was sealed with a gas permeable membrane and allowed to grow anaerobically for 24 h at 37 °C. Individual perturbations were prepared separately in the wells of a 96-deepwell plate. A single volume of the initial culture was prepared and transferred into each well to reduce the variability in initial populations between samples. In some experiments, replicate measurements were taken by recovering two cryo-stocks into cultures that were independently prepared and inoculated. For every set of experiments, multiple samples were devoted to a perturbation-free control. No tetracycline was included in the growth media on the day of the experiment.

Before preparing the growth perturbation experiment on day 1, at least six 1-mL aliquots of the original overnight culture were transferred into Eppendorf tubes and pelleted, the supernatant was removed, and the pellet was stored at −80 °C for future use as a control reference sample.

On the second day, the cultures were pelleted in the 96-deepwell plate, the supernatant was removed, and the cell pellets were stored at −80 °C for genomic DNA extraction.

### Fitness measurements from mouse colonization

All mouse experiments were conducted in accordance with the Administrative Panel on Laboratory Animal Care, Stanford University’s Institutional Animal Care and Use Committee. Experiments involved Swiss-Webster mice 6-12 weeks of age, and both genders were used.

Within four cages in a single germ-free isolator, five mice each were co-housed in cages 1–3 and sterilized water and food pellets were available *ad libitum*. The food available was a standard (Lab Diet JL Rand and Mouse/Auto 6F, 5K67*), MAC-deficient (Envigo Teklad custom diet TD.150689), or Western (Envigo Teklad custom diet TD.88137) diet in cages 1–3, respectively. Three mice were co-housed in cage 4 and fed a standard diet.

On day 0 of the colonization experiment, mice were orally gavaged with 200 µL of an overnight culture of the transposon pool grown anaerobically in commercial MRS with 10 µg/mL tetracycline. Mice in cage 4 were orally gavaged with 200 µL of 50 µM methotrexate in PBS on days 5, 6 and 7 of the experiment. Fecal pellets were collected daily for the first week and then every 2–3 days subsequently, and immediately stored at −80 °C. Two mice each from cages 1–3 were sacrificed on day 7 and sections of the gastrointestinal tract (ileum, cecum, and proximal colon) were stored at −80 °C. At the end of the experiment, all remaining mice were sacrificed and sections of their gastrointestinal tracts were stored. Mice were euthanized with CO_2_ and death was confirmed via cervical dislocation.

The mouse colonization experiment was repeated twice. In the first experiment, all mice were male, and the experiment was ended on day 5. In the second experiment, all mice were female, and the experiment was ended after one month.

### Fitness measurements from chick colonization

The germ-free chickens used in this study were housed at the Animal Research Welfare facility at South Dakota State University, US. The animal trial complied with the Institutional Animal Care and Use Committee guidelines and was conducted under approved protocol #18-058A.

Germ-free chickens were raised using a method adapted from our previously published protocol^86^. Briefly, Fertile Specific Pathogen Free (SPF) eggs were immersed in Sporicidin disinfectant solution (Contec 438, Inc.) for 30 s, then rinsed with sterile water to remove any sterilant residues. Eggs were aseptically transferred to a germ-free isolator and incubated at 37 °C with 55-70% humidity until hatching. Embryo-containing eggs were identified by candling.

One day post-hatching, chicks were randomly assigned to three groups, each containing 20 chicks. Germ-free status was confirmed by both anaerobic and aerobic culturing of pooled fecal material from each isolator. Chicks were given access to feed (Laboratory Chick Diet S-G 5065*, Lab Diet) and water *ad libitum*.

On the fifth day after hatching, chicks were orally administered two doses (12 h apart) of either a separate or combined culture of 10^8^ CFU/ml total of a either a *Bb* transposon pool, a transposon pool of *Bacteroides thetaiotaomicron*^20^ (*Bt*), or *Bb*+*Bt*. Subsequently, fecal samples were collected and stored at −80 °C. At intervals of 1, 2, 3, and 4 weeks post-inoculation, five chickens from each group were randomly selected for euthanasia by cervical dislocation. Aseptic collection of the contents and tissues from the small intestine, cecum, and colon were collected for further analysis.

### Analysis of fitness phenotypes

We used an established data analysis pipeline^18^ to analyze barcode abundance data and generate fitness metrics, with minor modifications. Using the established pipeline, we generated log_2_–fold change in abundance) values (*fitness*) to estimate effect size and a modified test statistic (*t*) to estimate significance.

In the initial quality control steps that determined which experiments were included in the dataset, we adjusted the cut-off of the quality control metric *adjcor* to be more permissive and accept more experiments, shifting the requirement from *adjco*r<0.25 to *adjcor*<0.35. We manually removed experimental conditions that visually produced little growth despite passing quality control metrics.

We next split experiments between *in vitro* perturbations and colonization time courses. *In vitro* experiments were averaged if they involved the same perturbation (condition and concentration). Fitness values were averaged using a mean. The. The test statistics were averaged based on whether the values were measured using the same cryo-stock (correlated barcode abundances in the control) or distinct cryo-stocks (separate barcode abundances in the control), as described previously^19^. Colonization experiments were averaged all samples from mice that shared the same cage and were collected on the same day. Experiments that were averaged from the colonization time course shared the same original cryo-stock and the test statistics were averaged accordingly (correlated barcode abundances in the control).

We defined statistically significant fitness phenotypes as having |*fitness*|>0.5 and |*t*|>4, and strong fitness phenotypes as having |*fitness*|>2 and |*t*|>5, as done previously^19^. To estimate the false discovery rate (FDR) of these cutoffs^19^, we compared the distribution of *fitness* and *t* values across all control samples to all experimental samples that passed our filtering steps and found that both definitions had FDR<5% (**Figure S1**).

When performing hierarchical clustering using fitness values, we found significant pairwise correlations within batches of samples performed on the same day and multiple genes with strong fitness defects across all conditions. These experimental artifacts raised the baseline correlations between experiments and genes and made correlation-based approaches like hierarchical clustering less statistically significant. To overcome this issue, we calculated a third statistic (*z*) from *fitness* and performed hierarchical clustering on this statistic. To calculate *z*, we first performed batch-wise normalizations of *fitness.* This step converts *fitness* from a direct measure of strain fitness to one of chemical interaction. Samples that were inappropriate for batch-wise normalization were filtered and, after filtering, the no-perturbation control experiments were removed from downstream analysis. Finally, the distribution of normalized fitness values were converted to a *z*-score for every gene. This *z* statistic displayed more significant correlations than *fitness* (**Figure S1**) and was used for defining statistically significant hierarchical clusters.

### Flow-sorting of progenitor collection

We used a previously developed protocol^21,80^ with minor modifications to isolate single strains from the transposon insertion pool. We confirmed that *Bb* did not produce H_2_S gas with exogenous L-cysteine added to the media, and thus included 10 mM L-cysteine in the MRS as an antioxidant during the sorting procedure^21^.

Commercial MRS was filter sterilized with a 0.2-µm PES filter and pre-reduced for 72 h. The day before the sorting procedure, filter-sterilized stocks were added to the MRS for a final concentration of 10 mM L-cysteine, 150 µM iron(II) sulfate heptahydrate, 100 µM calcium chloride dihydrate, and 10 µg/mL tetracycline hydrochloride. Two hundred microliters of this broth were aliquoted into the wells of 96-deepwell plates (Greiner, 786201) using an automated liquid handler (Mettler-Toledo, 30296707), and the 96-deepwell blocks were sealed with a foil seal (Thermo Scientific, 232698) and stored overnight at 37 °C in an anaerobic chamber.

The next morning, a cryo-stock of the transposon pool was diluted to an OD_600_ of 0.1 in pre-warmed MRS and recovered for at least 4 h at 37 °C. Over the course of the sorting procedure, this stock was repeatedly diluted to maintain a log-phase culture. An aliquot of the culture in a plastic test tube (Corning, CLS352003) and a batch of 30 96-deepwell plates were brought to the fluorescence activated cell sorter (FACS) (BD FACSJazz), and a restrictive gate based on FSC, SSC, and trigger pulse width was used to sort single cells into wells of the 96-deepwell plates. After inoculation in the FACS machine, the 96-deepwell plates were sealed with a gas permeable membrane, returned to the anaerobic chamber, and stored at 37 °C. 96-deepwell plates were transported to the anaerobic chamber in batches to minimize the time spent outside of the chamber, and the culture used for sorting was replaced every 45 min–1 h with a diluted log-phase culture from the anaerobic chamber. A final count of 172 96-deepwell plates were inoculated in this manner, collectively comprising the progenitor collection.

Cultures were recovered for 72-80 h, then 50 µL of filter-sterilized 50% (w/v) glycerol were added and the glycerol stock was aliquoted in triplicate in 96-microwell plates (Greiner, 651161). Plates were sealed with a foil seal and lid (Greiner, 656101) and stored at −80 °C.

### Identifying mutants for re-arraying from the progenitor to condensed collection

We used previously described protocols^21,80^ with minor modifications to locate mutant strains within the progenitor collection and choose optimal strains for re-arraying into the condensed collection. Notably, we used a plate-well pooling strategy^80^ that generated 268 (172+96) sequencing libraries. For inclusion in the condensed collection, we required that the barcode map to only one location in the genome (single-insertion strains) and we prioritized strains that only occurred once within the collection. When more than one insertion strain was found within the same gene, we prioritized the insertion closest to the middle of the gene body.

### Assembly of the condensed collection

We used a previously developed protocol^80^ with minor modifications to re-array select strains from the progenitor collection (172 96-well plates) into the final condensed collection (16 96-well plates). We used an EpMotion 5073 (Eppendorf) running EpBlue v. 40.5.3.10 inside of an anaerobic chamber for re-arraying.

Subsets of the progenitor collection were taken out of the −80 °C freezer and stored on dry ice. Lids were removed from the 96-microwell plates and cleaned with 70% (v/v) ethanol while the sealed 96-microwell plates were centrifuged (Fisherbrand, 14-955-300) for 30 s. After removing the seals, the sterilized lids were replaced and the 96-microwell plates were briefly returned to the dry ice bed for storage until they were stacked and transferred into the anaerobic chamber.

Forty microliters of thawed cryo-stocks were transferred from the 96-microwell plates of the progenitor collection to the condensed collection in 2-mL 96-deepwell plates (Greiner, 780270) previously inoculated with 1 mL of commercial MRS with 10 µg/mL tetracycline and pre-warmed at 37 °C. After transfer, the 96-microwell plates of the progenitor collection were resealed with a foil seal and returned to the −80 °C freezer. When fully inoculated, the 96-deepwell plates of the condensed collection were sealed with a gas-permeable membrane and transferred to a 37 °C incubator.

The inoculated cultures were incubated for 36–48 h at 37 °C. These cultures were used to inoculate fresh 96-microwell plates for growth curve analyses, then 250 µL of 50% (w/v) glycerol were added to each well and 18 copies of the condensed collection (60 µL aliquots) were transferred to new 96-microwell plates (Greiner, 651161). The plates were sealed and stored at −80 °C.

### Individual growth curve analyses of the condensed collection

Before adding 50% (w/v) glycerol to the 48-h cultures of the condensed collection, a small sample was used to inoculate fresh medium and OD_600_ was measured over time to generate growth curves, as in previous work^80^.

Briefly, 2 µL of the culture was transferred to 198 µL of fresh medium in a 96-microwell flat-bottom plate (Greiner, 655180) using an automated liquid handler. The media used for growth curves were commercial MRS, commercial RCM (Thermo Scientific, CM0149B), and commercial MRS with 300 mM sodium chloride. In some cases, 2 µL of an overnight culture of wild type were inoculated in an empty well as a positive control for growth. None of the media included tetracycline. The 96-microwell plates were sealed with plastic seals that had been cut so as not to extend over the edges of the plates to prevent plates from sticking to the stacker assembly. We used a plate stacker (BIOSTACK3WR, Biotek Instruments Inc.) and a Synergy H1 microplate reader (Biotek Instruments Inc.) running Gen 5 v. 3.08.01 to measure growth curves. The stacker and the front of the microplate reader were enclosed in a custom-fabricated box with a thermal control unit (AirTherm SMT, World Precision Instruments) to ensure a temperature of 37 °C. The plate stacker cycled through all plates every 30–42 min, which dictated the time between sequential OD readings. The plate stacker was disassembled after ∼48 h and the flat-bottom plates were removed from the anaerobic chamber for single-cell imaging.

### Single-cell imaging of the condensed collection

Stationary-phase cells from the growth curve cultures were diluted 1:10 into 0.85X phosphate buffered saline (PBS; Life Technologies, 70011-069) and transferred onto 1% agarose pads with 0.85X PBS to control for osmolality. Phase-contrast images were acquired with a Ti-E inverted microscope (Nikon Instruments) using a 100X (NA 1.40) oil immersion objective and a Neo 5.5 sCMOS camera (Andor Technology). Images were acquired using μManager v.1.4^87^. High-throughput imaging was accomplished using SLIP, as described previously^68^. Including sample preparation and calibration, SLIP enables acquisition of 49 images per well of a 96-well plate in ∼30 min.

### Determination of antibiotic minimum inhibitory concentrations

We measured changes in the minimum inhibitory concentration (MIC) for multiple genotoxic antibiotics in mutant backgrounds of *Bb* using a previously established approach^88^.

Briefly, individual insertion strains were isolated from the progenitor collection, single colony purified, and confirmed to carry the intended insertion using PCR. These strains were stored as independent cryo-stocks at −80 °C.

The strains were recovered overnight in commercial MRS with 10 µg/mL tetracycline, then diluted and grown into log phase. Log-phase cultures were inoculated at an initial OD_600_ of 0.01 in fresh commercial MRS in 96-microwell plates (Greiner, 655180) containing a dilution series of antibiotic at a final well volume of 200 µL. Prior to inoculation, we assembled the plate with only media and antibiotic and measured OD_600_ for use as a well-specific blank^89^. After inoculation, the plates were incubated at 37 °C and OD_600_ was measured every 1–5 min, depending on the experiment, to obtain growth curves. All OD_600_ measurements were obtained using an Epoch2 plate reader (Biotech) in an anaerobic chamber.

For each growth curve, we measured the area under the curve (AUC), defining the limits of integration between the initial OD_600_ measurement (*t_i_*) and 8 h past the point at which the growth rate (*µ*) fell below 10% of the maximum growth rate (*t_f_*). These limits were defined for each strain independently, using the no-antibiotic growth curve as a reference. AUC values were normalized against the no-antibiotic control (AUC_N_), and we defined the MIC for each strain as the first concentration at which AUC_N_ fell below 20%.

### Spot testing for CFU enumeration

We enumerated CFUs for dose-dependent killing of *Bb* by UV irradiation and the loss of viability of *Bb* upon exposure to glucose in a nutrient-poor environment. In both experiments, cultures were diluted 10-fold along the rows of a 96-microwell plate (Greiner, 651161). An automated liquid handler (Mettler Toledo) was used to transfer 0.5 µL of each dilution onto the surface of a rectangular Petri dish containing agar broth (MRS for the UV irradiation experiment and RCM for the glucose toxicity experiment). Spots were dried by leaving the Petri dish face up in an anaerobic chamber without a lid for ∼5 min, the lid was replaced, the Petri dishes were inverted, and the spots were incubated at 37 °C in the anaerobic chamber for 48 h. Colonies were counted at the dilution that had the highest concentration of cells yet crowding did not prevent accurate quantification.

For experiments involving UV irradiation (Figure 4**, Figure S4**), overnight cultures grown anaerobically in commercial MRS were used. For transposon insertion strains, selective pressure was maintained with 10 µg/ml tetracycline. After drying the spots, Petri dishes were removed from the anaerobic chamber and UV-irradiated with specific doses (mJ) in a UV Stratalinker 1800 (Stratagene) stored in a 37 °C room. Plates were then returned to the anaerobic chamber and incubated at 37 °C for 48 h. Pilot experiments showed that none of the strains tested were sensitized to 4 h of oxygen exposure (**Figure S4**), and the time spent outside of the chamber during UV irradiation was <1 h. As a control, we used a UVG-54 Handheld UV Lamp (Analytik Jena) housed in the anaerobic chamber to expose cells to UV irradiation for specific time periods (**Figure S4**). The results were qualitatively similar to irradiation outside the anaerobic chamber.

For experiments studying glucose toxicity (Figure 3), a time course of wildtype cultures grown in chemically defined media (CDM2) with various carbohydrates was used. Cultures were grown in 2-mL volumes in a 96-deepwell plate and sampled in a time course, and CFUs were enumerated at each time point.

### Fluctuation analyses

Cryo-stocks of single-colony, PCR-confirmed strains were inoculated into commercial MRS and grown overnight (∼14 h). Selective pressure for the transposon insertion strains was maintained using 5 µg/mL tetracycline. Overnight cultures were diluted 1:100 in fresh commercial MRS (no selective pressure) and grown at 37 °C until log phase at OD_600_ of 0.4–0.6 (∼5 h). Log-phase cultures of each strain were normalized to an OD_600_ of 0.2 and serially diluted in 10-fold steps to obtain a cell concentration of 10^3^ mL^-1^. This cell concentration was calculated based on a previously determined estimate of 5⨉10^8^ cells per OD unit of *Bb* during log-phase growth in commercial MRS.

Subsequently, 12 independent cultures were inoculated for each combination of strain and drug concentration by transferring 20 µL of culture dilution into 1 mL of fresh medium in a 2-mL 96-deepwell plate (for an initial cell concentration of 2×10^1^ mL^-1^). The concentrations of 4NQO used were 0 (untreated), 830 nM, and 6.2 µM. Initial experiments revealed that *Bb* has a strong inoculum effect^90^ with regards to 4NQO sensitivity and concentrations were empirically determined as half the MIC of 4NQO for the strains when inoculated at a cell concentration of ∼1×10^2^ mL^-1^. Wild type was grown in all three concentrations, while *scp3* and *uvrA* were not grown in 6.2 µM 4NQO as the treatment would have been lethal. All cultures were grown to early stationary phase, with an OD_600_ of 1–1.2 (∼24 h).

Two hundred microliters from each stationary-phase culture were spread on a pre-warmed commercial RCM agar Petri dish containing 5 µg/mL rifampicin. The plated cultures were incubated at 37 °C for 24 h in an anaerobic chamber and rifampicin resistant (*rif^R^*) colonies were enumerated.

The 4NQO-dependent mutation rate was estimated using the web application *bz– rates*^91^. Parameters input into *bz-rates* included the distribution of *rif^R^* CFUs, the estimate of cell counts from corrected OD_600_, and the fraction of the population plated (*z*=0.2).

### SOS-reporter quantification

An SOS-reporter was constructed using a transcriptional fusion of codon-adapted *msfGFP* and the 5’-UTR of the *Bb recA* open reading frame in the pNZ44 backbone (pNZ.*PrecA-msfGFP*), and transformed into the wild-type, *scp3* and *uvrA* genetic backgrounds.

Cryo-stocks of these reporter strains were recovered overnight in commercial MRS, with 5 µg/mL chloramphenicol and 10 µg/mL tetracycline for the transposon-insertion strains. These overnight cultures were diluted 1:100 into fresh MRS with 5 µg/mL chloramphenicol and grown into log phase. These log-phase cultures were used to inoculate a 96-microwell plate (Greiner, 655180655180) at an initial OD_600_ of .01. These final cultures were grown in commercial MRS with 5 µg/mL chloramphenicol and a dilution series of 4-NQO. We empirically determined the optimal concentration range, considering the inoculum effect of 4-NQO and the sensitization of strains to 4-NQO by the pNZ.*PrecA-msfGFP* plasmid. This 96-microwell plate was incubated at 37 °C in an Epoch2 plate reader (Biotech) and OD_600_ was monitored every 5 min. After the untreated control cultures entered early stationary phase (∼24 h), we removed the 96-microwell plate from the anaerobic chamber and incubated the cultures aerobically with agitation at 900 rpm (Eppendorf ThermoMixer FP) at 4 °C for 1 h. Cultures were then either directly spotted onto 1.5% (w/v) agarose pads (Invitrogen, 16500-500) with PBS, pH 7.4 (Life Technologies, 70011-069) or diluted into PBS, pH 7.4 to decrease cell density and then spotted onto PBS agarose pads.

Phase and fluorescence microscopy images were collected using a Ti-E microscope (Nikon) with a 100X (NA: 1.4) objective and a Zyla 5.5 sCMOS camera (Andor). Cells were segmented from phase-contrast images using *Morphometrics*^92^. The fluorescence signal was background-corrected by subtracting the median value of pixels in the fluorescence channel that were not within a cell contour, and quantified fluorescence as the median background-subtracted signal within each cell contour. We found that while the average fluorescence signal of the population did not substantially change in response to 4-NQO, a subpopulation exhibited more than a 10-fold increase in fluorescent signal around or above the 4-NQO MIC of wild type. We quantified this phenomenon using the 95^th^ percentile of fluorescence signal as a function 4-NQO concentration (Figure 4).

### Growth of strains for peptidoglycan analysis

Strains of interest were recovered from cryo-stocks via transfer into 2 mL of commercial MRS with 10 µg/mL tetracycline and grown anaerobically overnight at 37 °C. The next day, overnight cultures were diluted 1:10 into fresh medium, maintaining selection for the transposon insertion strains. Cells were kept in log phase via periodic dilution. Two-hundred fifty microliters of log-phase culture were used to inoculate 50 mL of medium without antibiotic in a conical centrifuge tube (Thermo Scientific, 339652).

Transposon insertion strains were inoculated in commercial MRS, and wild type was inoculated in both commercial MRS and MRS*. Cultures were grown anaerobically for 17 h, brought out of the anaerobic chamber, and stored on ice. All subsequent work was performed aerobically at 4 °C. Cultures were vortexed to fully suspend the cells, 1 mL was transferred to a 96-deepwell plate, and cells were pelleted from the remainder of the culture via centrifugation at 4,000 *g* and 4 °C for 15 min and stored at −80 °C. The 1-mL samples were used to measure OD_600,_ and cell shape and the cell pellets were used for cell wall compositional analysis.

Before analysis, each of the cell pellets was resuspended at a normalized OD_600_ by resuspension in ice-cold phosphate buffer saline (PBS; 137 mM NaCl, 2.7 mM KCl, 10 mM Na_2_HPO4, 1.8 mM KH_2_PO_4_, pH approx. 7.4) according to the pellet density factor, with the sample having the highest pellet density being resuspended in 50 mL of ice-cold PBS buffer. The OD_600_ was recorded before subsequent processing steps (Eppendorf D30 BioPhotometer).

### UPLC-MS analysis of peptidoglycan

Murein sacculi isolation and muropeptide analysis were essentially performed as described previously^93,94^. Four milliliters of each bacterial resuspension in ice-cold PBS were pelleted by centrifugation (14000 *g*, 10 min, 4 °C) (Heraeus Fresco 21, Thermo Fisher Scientific). The cells were then resuspended in 4 mL of PBS containing 2.5% sodium dodecyl sulfate (SDS). The mixture was boiled for 1 h and stirred for a further 20 h after the heat was turned off. SDS was removed by washing the sacculi four times with 2 mL of ultrapure water, pelleting was performed with a centrifuge (Heraeus Pico 21, Thermo Fisher Scientific; 21000 *g*, 15 min).

The sacculi pellets were resuspended in 1 mL of 100 mM Tris-HCl, pH 8. The samples were then treated with trypsin (Sigma-Aldrich T4799) by adding 50 μL of a 2 mg/mL stock solution and 25 μL of 50 mM CaCl_2_ and incubating for 1 h at 37 °C with agitation at 300 rpm in a Thermomixer comfort (Eppendorf, Germany). The cell wall material was pelleted and washed two times with 2 mL of ultrapure water (21000 *g*, 15 min).

To release the cell wall polysaccharides, the pellets were resuspended in 500 µl of 1 M HCl and incubated for 4 h at 37 °C with agitation at 300 rpm. The sacculi were then pelleted and washed five times with 2 mL of ultrapure water. Clean sacculi were resuspended in water and digested overnight with 100 µg/mL muramidase (in house preparation, final concentration) at 37 °C with agitation at 150 rpm. Reactions were inactivated by heating at 100°C for 5 min and centrifuged (21000 *g*, 15 min) to remove precipitated material.

Soluble muropeptides were reduced with 0.5 M sodium borate pH 9 and sodium borohydride at 10 mg/mL (final concentration) for 20 min at room temperature. The pH was adjusted to 3.5 by addition of orthophosphoric acid. The samples were finally centrifuged (5250 *g*, 10 min) in an Allegra X-15R centrifuge (Beckman Coulter, USA) using 96-well-filter plates (AcroPrep Advance 96-well 0.2 μm GHP 636 membrane filter plate, Pall, USA) with a collection plate (deep well storage plate, 96-well, 2,2 mL; Thermo Fisher Scientific, Germany).

Detection of muropeptides was performed on an Acquity H-Class UPLC coupled to a Xevo G2/XS QTOF mass spectrometer controlled with *MassLynx*^95^ software (version 4.2, Waters Corporation, USA). Chromatographic separation was achieved using a Kinetex C_18_ LC column (150 mm x 2.1 mm, 1.7 µm in particle size and 100 Å in pore size; Phenomenex, USA). The mobile phase consisted of a mixture of 0.1% formic acid in water (solvent A) or 0.1% formic acid in acetonitrile (solvent B) in an 18 min run. The solvent B gradient was set as follows: 0–3 min, 2%–5%; 3–6 min, 5%–6.8%; 6–7.5 min, 6.8%–9%; 7.5–9 min, 9%-14%; 9–11 min, 14%–20%; 11–12 min, hold at 20%; 12–12.1 min, 20%–90%; 12.1–13.5, hold at 90%; 13.5–13.6 min. 90%–2%; 13.6–18 min, hold at 2%. The gradient program was used with a flow rate as follows: 0–12.1 min, 0.25 mL/min; 12.1–16.1 min, 0.3 mL/min; 16.1–18 min, 0.25 mL/min. The temperature of the column and autosampler were maintained at 45 °C and 10 °C, respectively. The injection volume was 2 μL. UV detection was performed at 204 nm with a sampling rate of 20 points/s. The Xevo G2/XS QTOF was operated in sensitivity (ESI+) mode. The detection parameters of the ESI source were used as follows: capillary voltage, 3 kV; sampling cone, 20 V; source offset, 80 V; source temperature, 100 °C; flow rate of cone gas, 100 L/h; temperatures and flow rate of desolvation gas (N_2_), 300 °C and 500 L/h; and collision energy, 6 eV. Mass spectra were acquired at a speed of 0.05 s^-1^. Data dependent acquisition algorithm (FastDDA) was used as the mode of data acquisition. Spectra were first acquired from 100 to 2,000 *m/z* and then two precursors with the highest intensities were selected for fragmentation. Scan rate for MS/MS was 1 s. For the MS/MS, collision energy ramp was used for the fragmentation (Low Mass CE Ramp 20-40 V, High Mass CE Ramp 30-60 V). A compound library created in *ChemSketch*^96^ (version 2021.1.2 ACD/Labs, Canada) was used to detect and identify each muropeptide. Quantification was done by integrating peak areas from the extracted ion chromatograms (EICs) of the corresponding *m/z* value of each muropeptide using *TargetLynx XS* application manager (Waters Corp., USA).

The relative amount of each muropeptide was calculated by dividing the sum peak areas of considered ions of a muropeptide by the total extracted ion chromatograms. The abundance of peptidoglycan (total peptidoglycan) was assessed by normalizing the total ions peak area to the OD_600_. Monomers (%), dimers (%), and trimers (%) were calculated as sum of the relative abundances of all monomeric, dimeric or trimeric muropeptides. The degree of cross-linking was calculated as dimers + (trimers × 2).

### UPLC-MS analysis of intracellular muropeptide

Four milliliters of each resuspension were pelleted via centrifugation (14,000 *g*, 10 min, 4 °C) (Heraeus Fresco 21, Thermo Scientific), the cells washed 4 times with ice-cold 0.9% (w/v) sodium chloride, then the pellets were dissolved in 100 μL ultrapure water and boiled for 15 minutes. After clarification of the suspension by centrifugation at maximum speed in a benchtop centrifuge for 10 min (Heraeus Pico 21, Thermo Fisher Scientific), the supernatants were filtered through a 0.2-μm 96-well-filter plate (AcroPrep Advance 96-well 0.2 μm GHP 636 membrane filter plate, PALL) with a collection plate (96-deep well storage plate 2.2 mL; Thermo Fisher Scientific). The final protein concentrations in the samples were quantified using the Qubit Protein Assay Kit (Invitrogen, USA) and the Qubit 4 (Invitrogen, USA) and normalized by dilution in ultrapure water to a concentration of 0.1 µg/µL total protein.

Detection and characterization of peptidoglycan precursors by LC-MS was performed on an Acquity H-Class UPLC coupled to a Xevo G2/XS QTOF mass spectrometer controlled with *MassLynx*^95^ software (version 4.2, Waters Corporation, USA). Chromatographic separation was achieved using a Kinetex C_18_ LC column (50 mm x 2.1 mm, 1.7 µm in particle size and 100 Å in pore size; Phenomenex, USA). The mobile phase consisted of a mixture of 0.1% formic acid in water (solvent A) and 0.1% formic acid in acetonitrile (solvent B) in a 6 min run. The gradient of solvent B was as follows: 0–0.5 min, 2%; 0.5–3 min, 2%–10%; 3–3.5 min, 10%–20%; 3.5–3.8 min, 20%–80%; 3.8–4 min, hold at 80%; 4–4.1 min, 80%-2%; 4.1–4.6 min, 2%-80%; 4.6–4.7 min, hold at 80%; 4.7–4.8 min 80%–2%; 4.8–6 min, hold at 2%. The flow rate was 0.7 mL/min. The temperature of column and autosampler were maintained at 60 °C and 10 °C, respectively. The injection volume was 5 μL. UV detection was performed at 204 nm with the sampling rate of 20 s^-1^. The Xevo G2/XS QTOF was operated in sensitivity (ESI+) mode. Time-of-flight multiple reaction monitoring (ToF-MRM) was used as the mode of data acquisition. The transition list was created based on a compound library of expected precursors in *ChemSketch* (version 2021.1.2 ACD/Labs, Canada). The detection parameters of the ESI source were as follows: capillary voltage, 3 kV; sampling cone, 20 V; source offset, 80 V; source temperature, 100 °C; flow rate of cone gas, 100 L/h; temperatures and flow rate of desolvation gas (N_2_), 300 °C and 500 L/h; and collision energy, 6 eV. Mass spectra were acquired at a scan time of 0.036 s. The scan was in a range of 100–2000 *m/z*. The data was processed using the *TargetLynx XS* application manager (Waters Corp., USA).

The precursor abundances were quantified by integrating peak areas from extracted ion chromatograms (EICs) of the corresponding *m/z* value of each muropeptide precursor.

### Cell shape analysis

Phase-contrast images were first segmented with the deep learning-based software *DeepCell*^97^, using manually annotated representative images from the same image data set as the training dataset, and the resulting segmented images were analyzed using *Morphometrics*^92^ to obtain cell contours at sub-pixel resolution. Cell branches were inferred from the contours using the meshing algorithm in *Morphometrics* and filtered using custom MATLAB scripts to remove spurious branches. The length, width, and curvature of individual contours/branches were calculated based on the contour and branch information. >1500 cells were typically analyzed for each mutant.

### Spent medium metabolomics

Strains of interest were recovered from cryo-stocks, struck for single colonies on commercial MRS agar with 10 µg/mL tetracycline, and re-arrayed in the wells of a 96-deepwell plate. Cultures from single colony purification were grown for 48 h at 37 °C in 1 mL of commercial MRS with 10 µg/mL tetracycline. Cells were then pelleted via centrifugation at 4,000*g* x for 15 min at room temperature and washed three times using 900 µL of CDM1 with 1% glucose. Cells were resuspended in 900 µL of CDM1 with 1% glucose, and 64 µL of the resuspension were inoculated into 500 mL of CDM1 with 1% glucose. We estimated this culture to correspond to an initial OD_600_ of ∼0.2. The 96-deepwell plate was incubated at 37 °C for 48 h and supernatants were harvested for analysis. To recover the spent media, cultures were pelleted via centrifugation at 4,000*g* and 4 °C for 20 min, the supernatant was removed and aliquoted into polypropylene 96-well PCR plates (BioRad, AB3796), and the plates were sealed with foil seals (Thermo Scientific, 232698) for storage at −80 °C.

Samples were thawed only once, immediately before LC-MS/MS analysis. Samples were analyzed by two chromatography methods, reversed-phase (C18 column) and hydrophilic interaction chromatography (HILIC column). Protocol details and parameters are described in the Supplemental Information. Briefly, metabolites were extracted using extraction mixtures containing stable isotope-labeled internal standards. Samples for C18 analysis were dried at room temperature using a Labconco CentriVap, and reconstituted in 20% acetonitrile prior to analysis. Two microliters of prepared samples were injected onto a Waters Acquity UPLC BEH Amide column with an additional Waters Acquity VanGuard BEH Amide pre-column (HILIC) or Agilent SB-C18 column with a Phenomenex KrudKatcher Ultra filter frit attached to the column inlet (C18). The columns were coupled to a Thermo Vanquish UPLC machine. Chromatographic separation parameters^98^ and mass spectral parameters^99^ were described previously, with minor modifications. Spectra were collected using a Thermo Q Exactive HF Hybrid Quadrupole-Orbitrap mass spectrometer in both positive and negative mode ionization (separate injections, sequentially). Full MS-ddMS2 data were collected. Data were processed using *MS-DIAL* v. 4.60^100,101^. Alignment retention time and mass tolerance were set to 0.05 min and 0.015 Da, respectively. Aligned peaks were retained for further analyses only if they were present in at least two of three replicates and were >5-fold higher than the water blank average in at least one sample.

## Acknowledgements

We thank members of the Huang lab for helpful discussions, and Morgan Price for assistance with fitness analyses. This work was funded by a National Defense Science and Engineering Graduate Fellowship (to R.N.C.), Stanford Interdisciplinary Graduate Fellowships (to J.Sun and B.D.K.), NIH grants RM1 GM135102 (to A.L.S., K.C.H., A.M.D., J.Sonnenburg, and C.R.B.), R01 AI147023 (to K.C.H.), and R35 GM146949 (to. B.H.G.), NSF Awards EF-2125383 and IOS-2032985 (to K.C.H.) Science Foundation Ireland, through the Irish Government’s National Development Plan SFI/12/RC/2273-P1 and SFI/12/RC/2273-P2 (to L.F. and D.v.S.), the Swedish Research Council (to F.C. and M.N.), the Knut and Alice Wallenberg Foundation (to F.C. and M.N.), the Laboratory of Molecular Infection Medicine Sweden (to F.C. and M.N.), the Kempe Foundation (to F.C. and M.N.), a postdoctoral fellowship from Swedish Society for Medical Research (to M.N.), and USDA grants SD00R646-18 and SD00H702-20 (to J.Scaria). J.Sonnenburg, B.H.G., and K.C.H. are Chan Zuckerberg Biohub Investigators.

